# A machine learning framework for supervised treatment response prediction from tumor transcriptomics: A large-scale pan-cancer study

**DOI:** 10.1101/2025.10.24.684491

**Authors:** Lipika R. Pal, E. Michael Gertz, Nishanth Ulhas Nair, Sumit Mukherjee, Sumeet Patiyal, Thomas Cantore, Emma M. Campagnolo, Tian-Gen Chang, Saugato Rahman Dhruba, Yewon Kim, Eldad D. Shulman, Padma Sheila Rajagopal, Danh-Tai Hoang, Sridhar Hannenhalli, Alejandro A. Schäffer, Eytan Ruppin

## Abstract

Precision oncology aims to guide treatment decisions using biomarkers. While DNA-based panels are increasingly applied, RNA transcriptomics remain underused due to limited datasets and the absence of robust models. We assembled the largest transcriptomic resource for drug response prediction to date, spanning 91 cohorts, 5,675 patients, nine cancer types, and six frontline therapies: anti-PD-1/PD-L1 immune-checkpoint inhibitors, trastuzumab, bevacizumab, BRAF inhibitors, paclitaxel, and FAC/FEC (Fluorouracil-Adriamycin-Cyclophosphamide/Fluorouracil-Epirubicin-Cyclophosphamide) chemotherapy. We developed EXPRESSO (EXpression-Profile-RESponSe-Optimizer), a supervised machine-learning framework that predicts treatment response from pre-treatment transcriptomes by integrating drug targets and context-specific biomarkers. EXPRESSO achieves mean ROC-AUCs of 0.62–0.73 and median odds ratios of 2.4–4.6 across therapies, outperforming 20 published transcriptomic signatures and other machine learning methods. Prospective validation on 22 independent cohorts confirms that performance generalizes beyond cross-validation. The EXPRESSO signature additionally stratifies progression-free survival in immune checkpoint blockade-treated cohorts, demonstrating prognostic value beyond binary response prediction. Robustness analysis reveals that predictive performance plateaued for some therapies with increasing training cohorts but continued to improve for others. These findings suggest inherent limits of supervised brute-force learning for certain treatments, but additional data and deeper mechanistic modeling may further enhance transcriptomics-based predictors.

**SIGNIFICANCE:** EXPRESSO forms the next step in studying the feasibility of harnessing bulk transcriptomic data to inform therapeutic decision-making, advancing the role of transcriptomics from exploratory biomarker discovery to actionable predictive modeling.

## INTRODUCTION

Despite major advances in cancer therapeutics, not all patients benefit equally. Response rates vary widely, even within the same cancer type, due to underlying molecular heterogeneity. This challenge has fueled precision oncology, which aims to tailor therapies to individual patients using molecular biomarkers. To date, clinical implementation has largely focused on genomic profiling—particularly targeted DNA sequencing panels—to identify actionable mutations or fusions. However, since RNA alterations can reveal information beyond DNA(1), we set out to explore how genome-wide RNA expression can be leveraged to predict patient response to widely used cancer treatments on a large-scale.

Modern oncology now encompasses a broad array of therapeutic modalities, each targeting distinct aspects of tumor biology. These include immune checkpoint blockade, monoclonal antibodies (mAbs), targeted small molecule inhibitors, and traditional cytotoxic chemotherapies. Immune checkpoint inhibitors reinvigorate anti-tumor T cell responses by blocking inhibitory pathways (2). Monoclonal antibodies offer high specificity by binding to target antigens on cancer cells or the tumor microenvironment, often engaging immune-mediated mechanisms(3). Targeted small molecules inhibit key signaling proteins—typically kinases—associated with tumor-specific genetic alterations (4). Cytotoxic chemotherapies remain a mainstay for many cancers, exerting their effects through DNA damage or disruption of mitosis(5).

As new drugs continue to expand the therapeutic arsenal, the problem of which treatment regimen a cancer patient should receive has grown increasingly complex. Targeted tumor DNA sequencing has been transformative for subsets of patients (e.g., EGFR mutations in lung cancer, BRAF V600E in melanoma). Yet, many tumors lack actionable alterations, and identified mutations often fail to predict benefit across diverse contexts(6). Moreover, DNA-based features alone do not fully capture dynamic processes such as transcriptional adaptation, tumor–microenvironment interactions, or immune activity. These limitations have driven growing interest in RNA transcriptomics, which provides a functional, real-time readout of tumor biology through global gene expression patterns—capturing coordinated pathway shifts, cell states, and immune responses that may underlie treatment sensitivity(7).

Currently, most FDA approved cancer biomarkers for such therapies are genomic (e.g., EGFR, BRAF, BRCA1/2) or proteomic (e.g., HER2 amplification, PD-L1 protein expression)(8). For the 116 drugs with biomarkers listed on the OncoKB site (https://www.oncokb.org/oncology-therapies)(9), all biomarkers can be assessed by either DNA sequencing or protein assays (10). Although in some instances, RNA assays substitute for protein measurements, there are currently no AI-based mRNA gene signature biomarkers that integrate the expression levels of multiple genes simultaneously. In immunotherapy, high PD-L1 protein expression and genomic markers such as high tumor mutational burden (TMB) and microsatellite instability (MSI) are routinely used(11). However, these markers often lack pan-cancer applicability and may overlook broader transcriptomic contexts.

The best known FDA-approved prognostic transcriptomic tool is Oncotype DX, a multigene expression assay used in early-stage breast cancer to assess recurrence risk and to guide chemotherapy decisions(12). More broadly, transcriptomic predictors have demonstrated promising results across various therapeutic classes. T-cell–inflamed gene expression profiles, for example, have been associated with response to immune checkpoint inhibitors(13). Large-scale efforts, such as The Cancer Genome Atlas (TCGA), have further shown that expression-based models can, in certain contexts, outperform DNA-based features.

A substantial body of work has attempted to harness transcriptomic data through machine learning to build predictive models of drug response(14). Given the limited availability of large clinical datasets, many efforts have focused on training models using cancer cell lines and transferring models to make predictions to patient tumors. Pioneering studies predicted responses to agents like docetaxel, bortezomib, and erlotinib using cell line-derived gene expression models(15). Subsequent work extended this strategy by imputing drug response in patient tumors based on in vitro transcriptomic models(16). More recently, the Celligner algorithm mapped tumors to mRNA-expression-similar cell lines, revealing transcriptomic alignment often diverged from mutation similarity or methylation similarity(17).

Advanced deep learning strategies, including few-shot learning (18), context-aware models (CODE-AE) (19), interpretable frameworks (DrugCell) (20), integrative multi-omics approaches (21), and foundation model using concept-bottleneck architecture (22) have further expanded the landscape of transcriptome-informed prediction. Several efforts have also focused on combination therapy, using concepts like independent drug action (IDA) to predict synergistic effects from monotherapy data (23). Despite these innovations, none of these approaches for using transcriptomics to predict patient response has yet been translated into clinical practice.

Parallel to these computational advances, prospective clinical studies have begun to incorporate transcriptomic data into real-world decision-making. For example, Beaubier et al. (24) reported that RNA-seq and non-RNA immunotherapy-related biomarkers increased the proportion of patients matched to precision therapies from 29.6% to 43.4%. The WINTHER trial demonstrated the utility of transcriptomic-guided therapy selection across diverse cancers (25,26). Other studies, such as (27) and the TuPro initiative (28), have shown that multi-omics profiling, including RNA expression, can meaningfully influence clinical outcomes when integrated into tumor board recommendations.

Nevertheless, two major impediments continue to limit the clinical use of transcriptomic data: small sample sizes and poor harmonization between cross-platform expression data (29). With the emergence of expansive transcriptomic datasets—including expression profiles from responders and non-responders across a range of cancers—it is now feasible to systematically evaluate transcriptome-informed predictors (30,31).

In this study, we present a large-scale evaluation of the potential for pre-treatment transcriptomic data to predict patient response across a range of commonly used cancer therapies. To our knowledge, this represents the largest compilation of clinical trial-based transcriptomic datasets assembled for drug response prediction, enabling a systematic investigation of the predictive value of pre-treatment gene expression profiles. We developed EXPRESSO (<u>EX</u>pression <u>P</u>rofile <u>RES</u>pon<u>S</u>e <u>O</u>ptimizer), a biologically-grounded, machine-learning-based, supervised learning framework tailored to specific drugs and cohort settings. In two usage modes, EXPRESSO incorporates either the leading drug’s target gene(s) alone or in combination with selected biomarkers derived from differentially expressed genes between responders and non-responders. This framework serves to study the feasibility of harnessing bulk transcriptomic data to inform therapeutic decision-making.

## MATERIALS and METHODS

### Data collection and processing

We performed three systematic searches for suitable articles in multiple medical research databases: Embase [Nov, 2023] [RRID:SCR_001650], PubMed [Nov, 2023] [RRID:SCR_004846], and Trialtrove [Aug 5, 2023 data freeze, total 93,161 oncology clinical trials] databases. For the Embase and Trialtrove searches, we required that the study must be a (prospective) clinical trial, but we did not impose the clinical trial requirement in our PubMed search (**Supplementary Table S1A**). Our search criteria were designed to find patients’ pre-treatment bulk transcriptomic (RNAseq or microarray) data in studies where i) patients received targeted therapy (either monoclonal antibody or small molecule), immunotherapy, or only chemotherapy for a drug of interest and ii) the response data after the treatment are available (**Supplementary Table S2**). Studies were excluded if they lacked publicly available pre-treatment expression or patient response data; investigated cancer types outside the nine cancer types (melanoma, lung, breast, head and neck, gastric, mesothelioma, colorectal, colon, and rectal) analyzed in this study; investigated a frontline drug used in combination with another frontline drug, a drug not analyzed in this study, or belonged to a drug class represented by fewer than five cohorts; contained expression data from non-tumor sources (cell lines, organoids, PDX models, or blood); was funded by a commercial sponsor; had a cohort size of ≤10 patients; employed single-cell or spatial transcriptomic profiling; or exhibited a highly imbalanced responder/non-responder ratio. A complete list of screened studies with inclusion/exclusion status and reasoning is provided in **Supplementary Table S1B**. When a study was eligible for exclusion under more than one criterion, we tried to follow the order in the diagram, but if we had any doubt about an earlier reason and found a later reason to apply surely, then we chose the later reason. “Pre-treatment expression or response data not available” lumps together studies in which the necessary data were not collected at all and studies for which we could not obtain the necessary permissions in time to include the dataset. The full cohort selection process, including the number of studies excluded under each criterion, is summarized as a CONSORT-style diagram in **Supplementary Figure S1**.

To address potential technical batch effects arising from the use of mixed platforms (RNA-seq and microarray), we additionally applied frozen surrogate variable analysis (32) (fsva) for batch correction. As fsva did not improve model performance over rank normalization alone, rank normalization was retained as the primary harmonization approach for all downstream analyses. We required that each cohort partition the patients into two annotated two groups – responders and non-responders. Clinical trials with multiple arms (different treatment) were treated as separate cohorts. To identify the patient response labels (either via RECIST [Response Evaluation Criteria in Solid Tumors] or pCR [pathological Complete Response] labels), we followed the same criteria for responder/non-responder as mentioned in the original publications or in the metadata from dbGaP or GEO for each cohort.

For RNAseq data from GEO (33) (https://www.ncbi.nlm.nih.gov/geo/), we downloaded the RNAseq normalized counts TPM data directly. For microarray data from GEO, we also downloaded the data directly from GEO. While selecting genes across datasets, we required the gene symbols to be the same – we might miss some alias genes due to gene mapping issues. We used the collapseRows function from the WGCNA package (34) with the MaxMean method (probe with maximum value of mean expression across all samples) to resolve the duplicated genes with multiple probes in the microarray.

To analyze gene expression data from dbGaP (35), we first downloaded raw FASTQ files. Quality control (QC) of the raw sequencing reads was performed using FastQC [RRID:SCR_014583] to assess sequencing quality and detect potential issues. Low-quality reads and bases were filtered out using standard QC parameters such as read quality scores (Q30). The remaining high-quality reads were then processed using Salmon (36) with its default settings to estimate transcript-level TPM (Transcripts Per Million) and read counts. Subsequently, the tximport R package (37), with its default settings, was used to summarize transcript-level abundances from Salmon’s output into gene-level TPM and count matrices.

We used rank normalized (by patient across genes) expression data from each cohort for further downstream analyses.

### Sensitivity analysis of cohort size threshold

The default minimum cohort size threshold of ≥10 patients was selected to maximize the number of cohorts available for LOCOCV training set construction. To assess the robustness of performance estimates to this choice, a sensitivity analysis was conducted for drug cohorts containing at least some cohorts with fewer than 30 patients (ICB-melanoma, ICB-non-melanoma, trastuzumab, bevacizumab, and BRAFi). The threshold was varied from ≥10 to ≥30 patients in steps of 5, retaining only thresholds at which at least 5 cohorts qualified. In the original LOCOCV design, each cohort serves as the test set in turn, with all remaining cohorts forming the training set. For the sensitivity analysis, at each minimum size threshold, only AUCs from test cohorts meeting the size criterion were included in the summary statistics (mean AUC and median odds ratio); cohorts below the threshold were not removed from the training sets of other folds, and no models were retrained. This approach studies the effect of small test cohort size on reported performance estimates, without altering the underlying cross-validation design. Confidence intervals (95%) were obtained by bootstrap resampling at the cohort level, where cohort AUC values were resampled with replacement (2,000 iterations, same random seed), quantifying uncertainty in the cross-cohort mean AUC arising from between-cohort variability. Because the seed was intentionally reset to the same value at the start of each bootstrap call, thresholds retaining an identical set of cohorts produce the same mean AUC and confidence interval. Results are shown in **Supplementary Figures S2A-B**.

### Sensitivity analysis of sequencing platform

To evaluate whether the use of both RNA-seq and microarray platforms introduced a bias into performance estimates despite rank normalization, per-cohort ROC-AUC was compared by platform across all 91 cohorts (39 RNA-seq, 52 microarray), both unadjusted (Wilcoxon rank-sum test) and after adjusting for drug class (ordinary least squares regression, AUC ∼ Platform + Drug). To test whether any platform effect differed between cross-validation and generalization to newly, independently collected cohorts, a platform × evaluation-mode (LOCOCV vs. independent validation) interaction term was additionally fit in a more complex model. Results are shown in **Supplementary Figures S2C-D.**

### Clinical heterogeneity index calculation

#### Purpose

The clinical heterogeneity index (*H_w,d_*) was designed to provide a single, interpretable summary of how clinically heterogeneous – the collection of cohorts is for a given drug class. Because prediction generalizability is expected to degrade when cohorts span multiple cancer types, include many independent studies, or mix monotherapy with combination regimens, we sought a composite measure capturing all three dimensions simultaneously. This index is used descriptively — to contextualize why certain drug classes (e.g., bevacizumab) are more challenging for transcriptomic prediction than others (e.g., ICB-melanoma).

#### Construction

To compare the degree of heterogeneity across drugs, we computed a clinical heterogeneity index, *H*_w,d_ that integrates 3 sub-indices: cancer-typer diversity, cohort spread and treatment mix for a given drug.

#### Cancer-type diversity: *H_cancer_*_,*d*_

This sub-index checks for diversity of the cancer types.

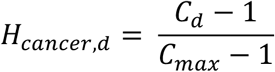

Where, *C_d_* is the number of distinct cancer types tested for drug d, and *C_max_* is the maximum number of types of cancer for any drug in the list. Linear normalization was chosen for simplicity, and this normalizes cancer-type spread between 0 (drug with single cancer type) and 1 (drug with most diverse cohorts).

#### Cancer-cohort balance index: *H_CC_*_,*d*_

This sub-index extends the cancer-type diversity score by incorporating the number of independent cohorts via a saturation function.

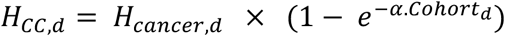

Where, 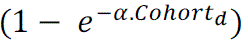 is a scaling function, where adding cohorts increases “balance” but with diminishing returns (the benefit of going from 1 to 2 cohorts is greater than going from 10 to 11 cohorts). Alpha (*α* = 0.15) is the saturation rate parameter, set empirically based on the range of cohort counts observed in our dataset, that controls how quickly the cohort correction approaches 1.

#### Treatment mix: *H_treatment_*_,*d*_

This sub-index checks for clinical variability.

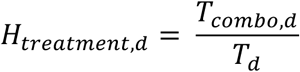

where, *T_combo_*_,*d*_ is the number of cohorts with combination treatment for a drug d and *T_d_* is the total number of cohorts for that drug d. Values near 1 indicate predominance of combination treatment (greater heterogeneity), while values near 0 indicate mostly monotherapy (greater homogeneity).

The final Heterogeneity index is defined in a weighted manner as:

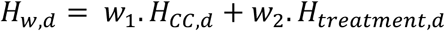

where the contribution of cancer-cohort balance for a drug d, w*_1_* = 0.4 and the contribution for treatment mix, w*_2_* = 0.6. We assigned greater weight (0.6) to the treatment mix component based on our intuition that variability among overall treatment regimen is more important than the cancer-cohort balance index.

### Generating LASSO Logistic Regression Models

We utilized the glmnet R package (38) to create LASSO logistic regression models from the rank-normalized features (details in **Supplementary Materials S1**).

### Leave-One-Cohort-Out Cross-Validation (LOCOCV) Framework

To ensure robust and generalizable evaluation, we employed a Leave-One-Cohort-Out Cross-Validation (LOCOCV) framework for the supervised learning (details in **Supplementary Materials S1**).

### EXPRESSO

EXPRESSO is a drug-specific machine-learning-based supervised learning framework specifically tailored for transcriptome-based drug response prediction. EXPRESSO leverages LASSO regression with L1 regularization, chosen for its capability to perform automatic feature selection and mitigate overfitting. EXPRESSO incorporates the entire transcriptome as input for feature selection and model training. It further integrates biological knowledge in the model using two strategies: (a) keeping the target gene(s) only as penalty-free (EXPRESSO-T, **Supplementary Materials S1**), or (b) using additional context-specific genes together with the target gene(s) as penalty-free (EXPRESSO-B, **Supplementary Materials S1**). Biomarkers were identified in a drug-specific manner using nested LOCOCV framework (**Supplementary Materials S1**). For downstream analyses, we retained only the variant with the higher mean ROC-AUC between the two modes. We used two baseline models – a supervised ‘vanilla’ model without any biological priors and an unsupervised target only model to compare the performance of EXPRESSO (**Supplementary Materials S1**).

### ICB-lung model

To evaluate cancer-type-specific modeling within the ICB-non-melanoma category, a dedicated EXPRESSO model was trained using the five NSCLC cohorts from the ICB-non-melanoma dataset. Model training, LOCOCV evaluation, and feature selection followed identical procedures to the primary EXPRESSO models (see LASSO Logistic Regression Models and EXPRESSO-B). Prospective validation was performed on a held-out NSCLC cohort (GSE274975) not used during model development, using both pooled and fold-averaged evaluation strategies as described under Independent Validation.

### Performance measures

Model performance in each fold of the LOCOCV framework was evaluated using both discrimination and effect size metrics. The primary metric was the area under the ROC curve (ROC-AUC), calculated from predicted probabilities on the held-out cohort. We also computed odds ratios (OR) by comparing response rates between patients predicted positive (score ≥ 0.5) and the remainder of the cohort, with 95% confidence intervals from Fisher’s exact test. For each drug, mean ROC-AUC and median OR across folds were reported, with drug-class–level summaries based on median OR to reduce outlier effects. Statistical comparisons between models used paired Wilcoxon signed-rank tests and DeLong’s test, where applicable (details in **Supplementary Materials S1**).

### Independent Validation

For four drug categories with available external data, we performed prospective-style independent validation using cohorts not used during model development. A single pooled model was trained on all original cohorts combined, and each independent cohort was tested against this model. As a sensitivity analysis, we additionally evaluated a fold-averaged approach in which each independent cohort was tested against every LOCOCV fold model (each trained with one original cohort left out) and the per-cohort ROC-AUC values were averaged across folds. For chemo-FAC-FEC, where sufficient independent cohorts were available (n=15), we benchmarked EXPRESSO against 20 published transcriptomic signatures by applying each method to the same held-out cohorts and comparing mean ROC-AUC.

### Survival Analysis

Cox proportional hazards regression and Kaplan-Meier survival analyses were performed for cohorts with available time-to-event data. The EXPRESSO signature score from each LOCOCV fold was used as a continuous predictor of PFS or OS. Survival times were standardized to months. Kaplan-Meier group stratification used an optimal cutpoint determined by maximizing the log-rank statistic (surv_cutpoint, survminer R package (39)). Cox model significance was assessed by the Wald test and Kaplan-Meier group differences were assessed by the log-rank test.

### Incremental Training for Upper-Bound Performance Estimation

To estimate the upper bound of predictive accuracy and assess the effect of training cohort size, we performed incremental training within the LOCOCV framework. For each test cohort, models were retrained iteratively by adding one training cohort at a time (from a single cohort up to all available cohorts, excluding the test cohort) and evaluated on the fixed test cohort. This stepwise accumulation allowed us to track how ROC-AUC changed with increasing training data and to approximate performance saturation points. To further evaluate scaling, we also plotted ROC-AUC against the number of training patients (rather than cohorts) across runs (**Supplementary Figures S3A-G**) (details in **Supplementary Materials S1**).

## DATA/CODE AVAILABILITY

All datasets are publicly available, but some require dbGaP permission (**Supplementary Table S2**). The code is available on Zenodo [RRID:SCR_004129] (https://doi.org/10.5281/zenodo.21878615) and in GitHub (https://github.com/ruppinlab/EXPRESSO). We provide the publicly available cohorts as a MultiAssayExperiment object.

## RESULTS

### Study overview

To assess the predictive power of bulk transcriptomic data in cancer therapy response, we aggregated publicly available (some requiring dbGaP permission) pre-treatment bulk transcriptomic cohorts from cancer clinical trials. Cohort selection followed eleven pre-defined exclusion criteria (**Methods**; **Supplementary Figure S1**; **Supplementary Table S1B**) applied to 584 candidate studies identified across three databases. The leading reasons for exclusion were cancer types outside the nine analyzed in this study, unavailable pre-treatment expression or response data, and frontline-drug-related criteria (combination therapy, a drug not analyzed in this study, or a drug class represented by fewer than five cohorts). An initial search identified 45 publications that met all inclusion criteria, yielding 69 cohorts (reflecting several publications that each contributed multiple cohorts of varying treatment arms) comprising 3,729 patients; during the manuscript revision, a second search of 340 additional studies using the same criteria identified 20 further publications, yielding 22 independent validation cohorts. Together, the full collection spans 91 cohorts, 5,675 patients, nine cancer types, and four major therapeutic modalities: immune-checkpoint-blockade, monoclonal antibody targeted therapy, small molecule targeted therapy, and chemotherapy (**Figure 1A, Supplementary Table S2, Supplementary Figure S1**). These cohorts included patients treated with six drug classes: anti-PD-1 checkpoint blockade (targeting the PD-1/PD-L1 axis; melanoma: 9 cohorts, 336 samples [initial] + 1 cohort, 30 samples [validation]; non-melanoma: 14 cohorts, 607 samples [initial] + 3 cohorts, 81 samples [validation]), trastuzumab (anti-HER2 monoclonal antibody; 13 cohorts, 494 samples [initial] + 3 cohorts, 141 samples [validation]), bevacizumab (8 cohorts, 377 samples), BRAF inhibitors (8 cohorts, 193 samples), paclitaxel (9 cohorts, 933 samples), and chemo-FAC-FEC (Fluorouracil-Adriamycin-Cyclophosphamide / Fluorouracil-Epirubicin-Cyclophosphamide; 9 cohorts, 790 samples [initial] + 15 cohorts, 1,694 samples [validation]) (**Figure 1B**). While most of the patients received these drugs in combination with varying chemotherapies and other drugs – as shown for the initial cohorts in **Figure 1C** and **Supplementary Table S2** — our prediction efforts focused solely on response to the six leading drug classes. All cohorts underwent consistent pre-processing and normalization to ensure data uniformity. Expression data from different platforms were rank normalized before downstream analyses (**Methods**).

**Figure 1:**
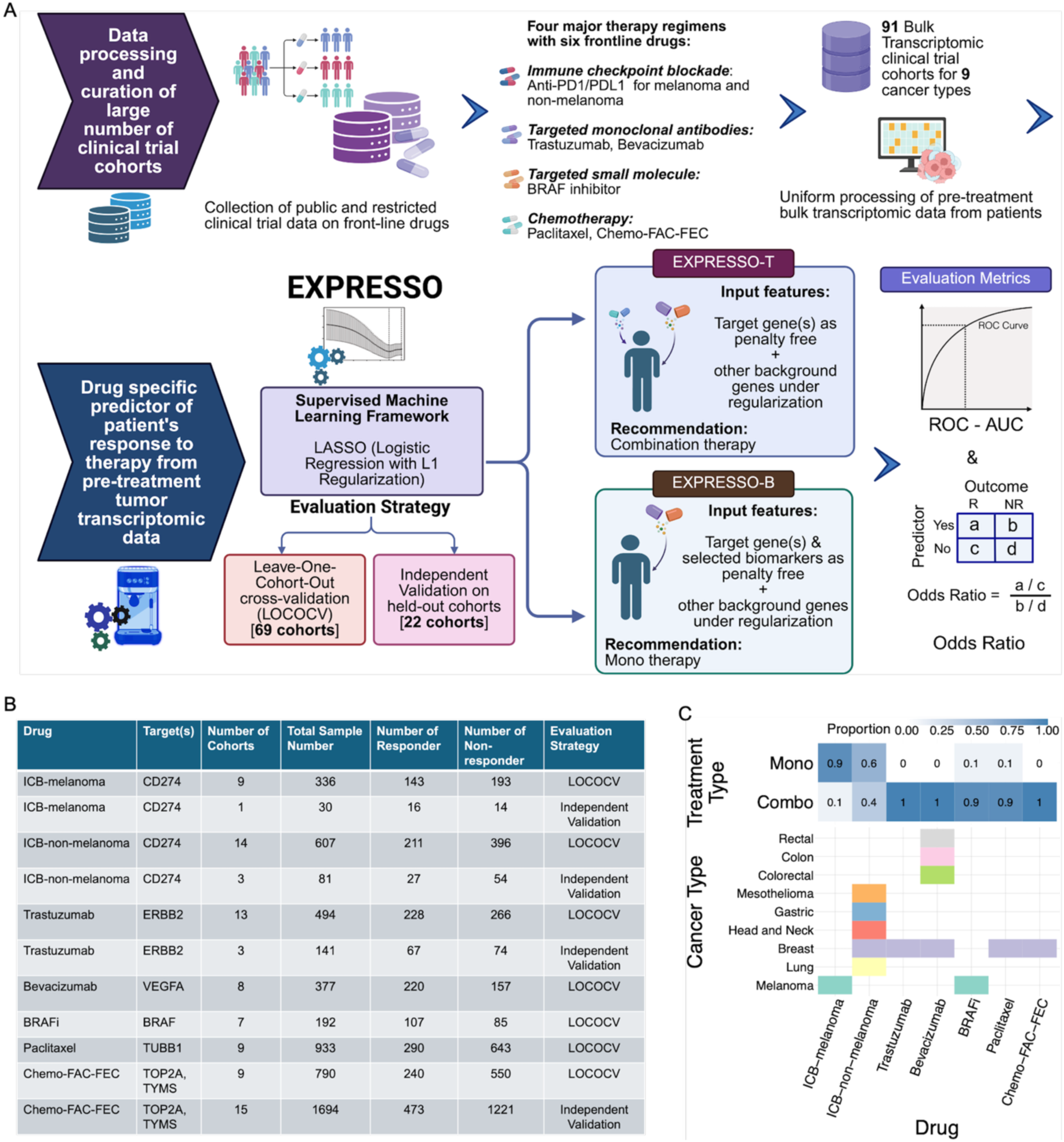
Study Overview and EXPRESSO workflow. (A) The workflow illustrating the evaluation of pre-treatment transcriptomic data for predicting patient response to cancer therapy. The analysis consists of two main components — first, data processing and curation of a large number of clinical trial cohorts, and second, the development of EXPRESSO — a machine-learning-based, biologically grounded, drug-specific predictor of therapy response from pre-treatment tumor transcriptomes (created in BioRender). In total, 91 bulk transcriptomic clinical trial cohorts spanning 9 cancer types were uniformly processed. EXPRESSO integrates two variants of biological information as input features in its LASSO (Logistic Regression with L1 Regularization) model: EXPRESSO-T, which uses only the drug’s known target gene(s) as a penalty-free feature, and EXPRESSO-B, which augments this setup by also including context-specific biomarkers. Model performance was evaluated using two strategies: Leave-One-Cohort-Out Cross-Validation (LOCOCV) across 69 cohorts, and independent validation on 22 held-out cohorts. Performance was assessed using ROC-AUC and odds ratio. Based on our study findings, EXPRESSO-T is recommended for combination therapy settings, whereas EXPRESSO-B performs optimally for monotherapy. (B) Summary table of the clinical trial cohorts used in this study, including the number of cohorts, total sample size, number of responders and non-responders, and evaluation strategy (LOCOCV or independent validation) for each drug class. (C) Characteristics of different drug classes in terms of treatment type (either mono or combination therapy, the darker blue corresponds to higher fraction of cohorts treated with combination therapy and the lighter blue corresponds to the fraction of cohorts with monotherapy) and cancer types of the patients in each drug class. Color codes for different cancer types are: melanoma – sea green, lung – yellow, breast – lavender, head and neck – red, gastric – steel blue, mesothelioma – orange, colorectal – light green, colon – pink, and rectal – gray.

To characterize transcriptomic variability among cohorts treated receiving the same drug class, we performed principal components analysis (PCA) on the rank-normalized pre-treatment gene expression data. This analysis revealed substantial variability across cohorts receiving the similar treatment with the same drug class (**Supplementary Figure S4**). To quantify the degree of clinical heterogeneity for each drug class, we employed a weighted index (H*_w,d_*) that integrates cancer-type diversity, cohort spread, and treatment composition (**Materials and Methods**, **Supplementary Table S3**). Among all treatment groups, the bevacizumab cohorts exhibited the highest heterogeneity (H*_w,d_* = 0.81), whereas ICB-melanoma cohorts showed the lowest heterogeneity (H*_w,d_* = 0.07).

Using the uniformly processed rank-normalized expression data, we first tested whether expression levels of known drug targets alone could predict treatment response, without any model training. We then developed EXPRESSO, a drug-specific supervised LASSO (L1-regularized) logistic regression framework that leverages the full transcriptome while incorporating biological priors. EXPRESSO integrates two variants of biological information as input features in its LASSO model: EXPRESSO-T, where only the drug’s known target gene is included as a penalty-free feature, and EXPRESSO-B, which augments this setup by also including context-specific biomarkers identified through a nested discovery procedure (**Supplementary Materials S1**). Known drug targets included *CD274* (anti–PD-1 inhibitors; separate models for ICB-melanoma and ICB-non-melanoma), *ERBB2* (trastuzumab), *VEGFA* (bevacizumab), *BRAF* (BRAF inhibitors), *TUBB1* (paclitaxel), and *TOP2A* and *TYMS* (FAC/FEC chemotherapy) (**Figure 1B**). As controls, we built two baseline models: (i) a ‘vanilla’ supervised LASSO model without drug target(s) or biomarker priors, to assess the added value of biological information, and (ii) an unsupervised target-only model. To mimic real-world applications and minimize overfitting, we used a Leave-One-Cohort-Out Cross-Validation (LOCOCV) framework. Model performance was assessed using ROC-AUC and odds ratios (**Methods**).

### Target expression alone demonstrates meaningful predictive power only in select monoclonal antibody, immunotherapy, and targeted therapies

We assessed whether expression of the drug target gene(s) alone (unsupervised mode) could predict treatment response across the seven sets of treatment-type cohorts (**Figures 2A-G**). Patients within each cohort were ranked based on the rank-normalized expression of each drug’s target gene, under the hypothesis that its expression might correlate with response. Across all drugs, the predictive performance of target gene expression alone varied widely, with mean ROC-AUCs ranging from 0.47 to 0.69. These findings suggest that while target gene expression may carry some predictive signal, it is insufficient for robust prediction.

**Figure 2:**
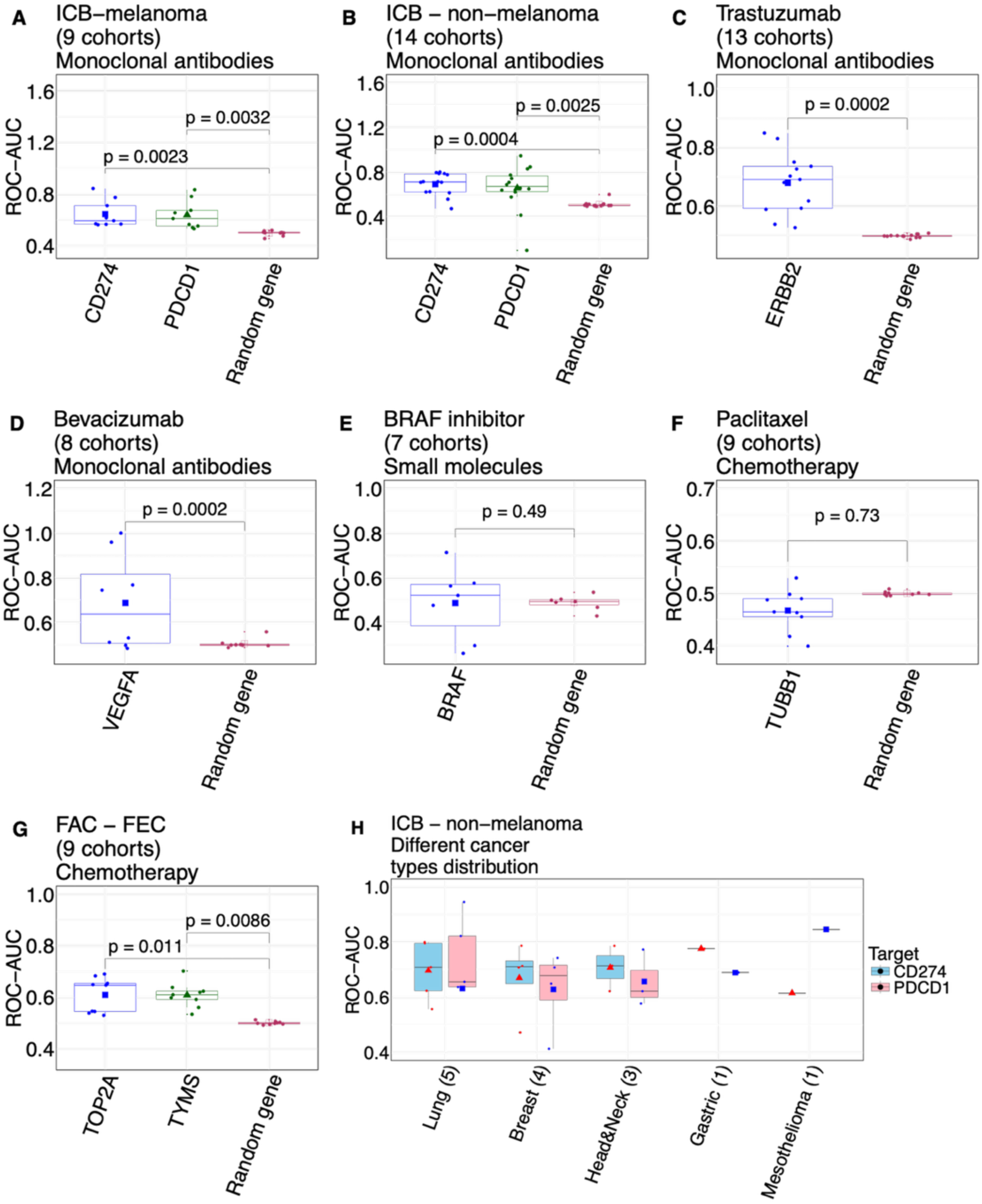
Drug response prediction based on target gene expression in an unsupervised approach. (A-G) Box plots showing ROC-AUC scores of the expression of target genes and median ROC-AUCs of corresponding random controls for all six drug classes. For immunotherapy, we separated melanoma and non-melanoma cohorts. Empirical p-values comparing target expression with random controls (10,000 randomly selected genes) are shown. (H) Box plots showing ROC-AUC scores of the expression of target genes for ICB-non-melanoma for different cancer types.

Notably, monoclonal antibody therapies—whether targeted or immunotherapeutic—showed a meaningful association between target gene expression and clinical response. For anti–PD-1 immune checkpoint blockade (ICB) therapies, *CD274* (encodes PD-L1), expression showed moderate predictive power across 9 melanoma cohorts (mean ROC-AUC 0.65, empirical p-value = 0.0023), with *PDCD1* (PD-1) expression performing similarly (mean ROC-AUC 0.64; **Figure 2A**). In 14 non-melanoma ICB cohorts—including lung, gastric, breast, head and neck, and mesothelioma—*CD274* expression again predicted response modestly well (mean ROC-AUC 0.69, empirical p-value = 0.0004; **Figure 2B, 2H**), slightly outperforming *PDCD1* (mean ROC-AUC 0.66), consistent with prior reports (40). For trastuzumab-treated breast cancer patients, higher *ERBB2* expression predicted therapeutic response, achieving a mean ROC-AUC of 0.68 (empirical p-value = 0.0002) across 13 cohorts (**Figure 2C**). Similarly, *VEGFA* expression modestly predicted response to bevacizumab in 8 cohorts across triple-negative breast, colorectal, rectal, and colon cancers, also yielding a mean ROC-AUC of 0.68 (empirical p-value = 0.0002; **Figure 2D**).

In contrast, *BRAF* expression did not predict response to BRAF inhibitors in 7 melanoma cohorts (mean ROC-AUC 0.49, empirical p-value = 0.49; **Figure 2E**). This is not surprising because *BRAF* inhibitors target an aberrant protein with a p.V600 mutation (41). Likewise, TUBB1, the known target of paclitaxel (42), showed no predictive power across 9 breast cancer cohorts (mean ROC-AUC 0.47, empirical p-value = 0.73; **Figure 2F**). For FAC-FEC chemotherapy, the expression of TOP2A (primary target of anthracyclines, like doxorubicin and epirubicin) (43) and TYMS (primary target of 5-fluorouracil) (44) showed moderate predictive performance (mean ROC-AUCs of 0.61 separately for each single gene across 9 cohorts, empirical p-value = 0.011 for TOP2A and empirical p-value = 0.0086 for TYMS; **Figure 2G**). Empirical p-values are based on control analyses using 10,000 randomly selected genes (**Methods**).

Together, these findings reveal that monoclonal antibody efficacy is more tightly linked to target gene expression, possibly due to the super-linear dependence (45) of antibody binding on target abundance. However, target expression alone is generally insufficient for accurate patient stratification. To go beyond target gene expression, we next opted for whole transcriptome analysis rather than limited gene panels, selecting genes measured in at least three cohorts for any specific treatment type to ensure comparability across datasets.

### EXPRESSO effectively predicts patients’ treatment responses for seven different drug regimens

We developed seven drug-specific, biologically informed, EXPRESSO LASSO models using whole-transcriptome features with regularization. Biologically informed priors were incorporated either as target gene(s) alone (EXPRESSO-T) or combined with differentially expressed (DE) biomarkers from the training data (EXPRESSO-B, **Methods**), both treated as penalty-free features. Model performance was evaluated using a LOCOCV strategy. Results were robust to the choice of minimum cohort size of ten samples (**Supplementary Figures S2A-B**) and did not differ significantly by sequencing platform, whether assessed across all cohorts, within LOCOCV alone, after adjusting for drug class, or when testing for a platform-by-generalization interaction (all p≥0.14; **Supplementary Figures S2C-D**).

Overall, the two EXPRESSO variants achieved comparable performance where available (**Supplementary Figures S5A-B**), except in ICB-melanoma, where EXPRESSO-B significantly outperformed EXPRESSO-T (p = 0.02, one-sided DeLong’s test). Notably, ICB-melanoma exhibited the lowest clinical heterogeneity index (H*_w,d_* = 0.07) among all drugs (**Supplementary Table S3**). In this setting, EXPRESSO-B identified additional biologically relevant genes – such as *IRF1* (46) and *UBD* (47) – beyond EXPRESSO-T features, thereby enhancing prediction accuracy ( **Supplementary Materials S2, Supplementary Figures S6A-Q**). For downstream analyses, we retained only the variant with the higher mean ROC-AUC between the two modes.

The variability in per-cohort ROC-AUC across LOCOCV folds reflects multiple sources of inter-cohort heterogeneity, including differences in response definitions (RECIST vs. pCR), sequencing platforms (RNA-seq vs. microarray), cancer-type composition, and combination treatment regimens. These factors are captured in part by our clinical heterogeneity index (H*_w,d_*), with higher-heterogeneity drug classes (e.g., bevacizumab, H*_w,d_* = 0.81) exhibiting broader AUC distributions than lower-heterogeneity classes (e.g., ICB-melanoma, H*_w,d_* = 0.07). Rank normalization was applied as a platform-agnostic harmonization strategy; standard batch correction approaches did not improve performance and were not retained.

Across all seven drug-specific models, EXPRESSO achieved mean ROC-AUC values ranging from 0.62 for BRAF inhibitors to 0.73 for trastuzumab (**Figures 3A-G**) and median odds ratios (ORs) from 2.4 for bevacizumab to 4.6 for ICB-melanoma (**Figure 3H**). While both metrics reflect model performance, they capture distinct aspects of predictability and are not expected to rank drugs identically. ROC-AUC is a threshold-independent measure that quantifies the model’s overall ability to discriminate responders from non-responders across all possible decision thresholds. The OR, by contrast, is a threshold-dependent effect-size measure that quantifies the enrichment of true responders among patients classified as predicted-positive at a fixed score cutoff (≥0.5). Consequently, divergences between the two metrics — such as the relatively modest ROC-AUC but high OR observed for ICB-melanoma — are expected and informative: they indicate a setting where the model confidently identifies a high-confidence subset of responders even if global discrimination across the full score range is moderate. Together, the two metrics provide complementary and more complete characterization of predictive performance than either alone. Compared to two baselines—a supervised vanilla model without biological priors and an unsupervised target-only model – the biologically informed EXPRESSO models consistently outperformed both (**Figure 3**). For bevacizumab, although the mean ROC-AUCs for EXPRESSO-T and the unsupervised target only model were identical (0.69), EXPRESSO-T exhibited markedly lower inter-cohort variability and significantly higher paired within-cohort ROC-AUCs (one-sided DeLong’s p-value = 0.03; **Figure 3D**), reflecting improved consistency across datasets. Performance gains were most pronounced for targeted and immunotherapies, whereas chemotherapy models showed minimal improvement, reflecting limited predictive signal from target genes.

**Figure 3:**
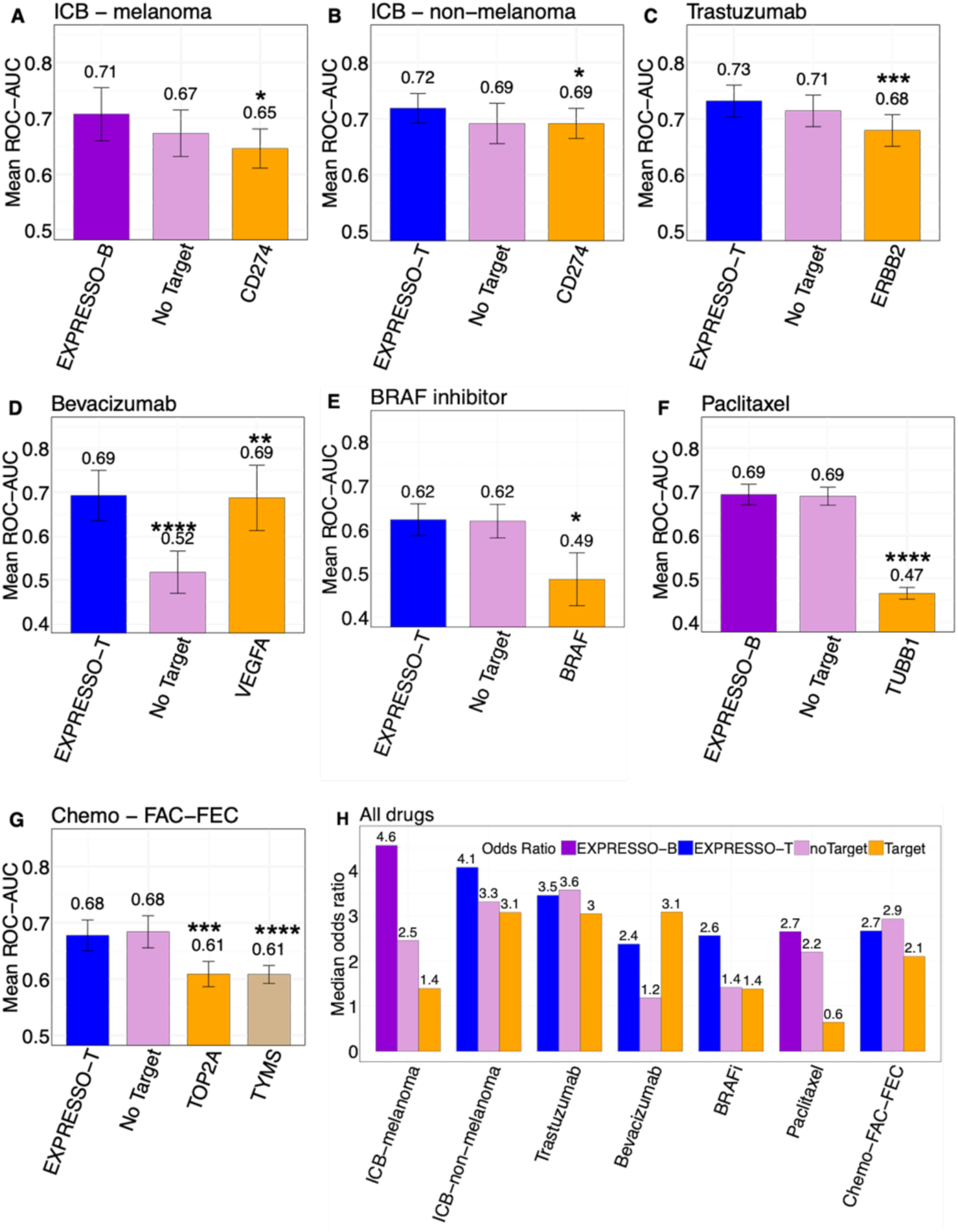
Drug response prediction of EXPRESSO compared with the two comparator baseline models. (A–G) Bar plots of mean ROC-AUC with standard error bars for all seven drug-specific EXPRESSO models (EXPRESSO-T in blue and EXPRESSO-B in dark violet), vanilla supervised models without biological priors (No Target, plum), and unsupervised target-only predictors (orange for the first target and tan for the second target). The best-performing EXPRESSO variant (by higher mean ROC-AUC between EXPRESSO-T and EXPRESSO-B) is shown for each drug, as the two EXPRESSO variants achieved comparable performances (bevacizumab and BRAFi have no EXPRESSO-B model), except ICB-melanoma (**Supplementary Figure S5**). Significant p-values (one-sided DeLong’s test across all cohorts for a drug) between EXPRESSO and no target supervised model or between EXPRESSO and unsupervised target model are denoted by an asterix (*) with significance levels as follows: * p ≤ 0.1, ** p ≤ 0.05, *** p ≤ 0.01, **** p ≤ 0.001. (H) Median odds ratios for the seven drug-specific EXPRESSO models are shown in the bar plot, using the same color scheme as the preceding figure. For Chemo-FAC-FEC, the reported value for the target represents the median across all cohorts predicted by the two target genes.

### EXPRESSO as the method of choice for clinical transcriptomic drug response prediction

To confirm that EXPRESSO’s modeling choices are well-suited to this prediction problem, we benchmarked EXPRESSO against seven ML methods (Random Forest, XGBoost, SVM-RBF, SVM-Linear, KNN, Neural Network, deep MLP) under identical LOCOCV conditions across seven drugs (**Supplementary Figures S7A-G**). EXPRESSO achieved the highest or joint-highest mean AUC for five out of seven drugs (ICB-melanoma: 0.71, ICB-non-melanoma: 0.72, trastuzumab: 0.73, paclitaxel: 0.69, Chemo-FAC-FEC: 0.68), with no nonlinear method significantly outperforming EXPRESSO in any drug (two-sided Wilcoxon p > 0.05). KNN performed significantly worse than EXPRESSO in five drugs. These results confirm that drug response prediction in our transcriptomic dataset is well-captured by a linear model, and that EXPRESSO’s regularization and interpretability advantages are not traded off against predictive performance.

### EXPRESSO-selected features capture drug-relevant biological pathways and immune landscape

Having established that EXPRESSO outperforms both baseline models and alternative machine learning methods, we next examined whether its predictive features are biologically coherent and whether its predictions reflect genuine transcriptomic signal beyond immune cell infiltration.

Reactome pathway enrichment analysis of stable feature sets identified significant pathway enrichment in four models (**Supplementary Figure S8, Supplementary Table S4**). Notably, the ICB lung model was enriched for complement cascade and inflammasome pathways, consistent with the innate immune character of its stable feature set, while the ICB non-melanoma model showed enrichment for G1/S cell cycle progression and TGF-β/SMAD signaling pathways. RHO GTPase cycle pathways were enriched in ICB melanoma. For the remaining models, the small number of genes identified precluded formal enrichment analysis; biological interpretability was assessed through manual feature annotation as described above.

To assess whether EXPRESSO predictions are confounded by tumor immune infiltration, we ran CIBERSORT (48) deconvolution on all ICB cohorts and correlated sample-level EXPRESSO prediction scores with estimated cell-type fractions. EXPRESSO scores showed moderate correlation with estimated CD8^+^ T-cell fraction (ICB-melanoma: Spearman r = 0.46, Benjamini-Hochberg (BH)-adjusted p < 0.001; ICB-non-melanoma: r = 0.15, BH-adjusted p = 0.002), but the strongest and most consistent correlation across both cohort sets was with M1 macrophage fraction (r = 0.47 and r = 0.33 respectively), accompanied by negative correlations with M2 macrophages (r = −0.16 and r = −0.17) and M0 macrophages (r = −0.19 and r = −0.17). This pattern reflects IFN-γ-driven macrophage polarization rather than lymphocyte infiltration alone, suggesting that EXPRESSO captures a broader immune activation signature.

In a direct comparison against simpler immune scores, EXPRESSO achieved the highest mean per-cohort AUC in both cohort sets compared to TIS, *CD8A* expression, and ESTIMATE immune score (**Supplementary Table S5**). In the ICB-non-melanoma cohort set, EXPRESSO significantly outperformed TIS (EXPRESSO wins in 10/14 cohorts, paired Wilcoxon p = 0.045) and ESTIMATE immune score (10/14 cohorts, p = 0.026). Comparisons in the smaller ICB-melanoma cohort set (n=9 cohorts) were in the same direction but did not reach statistical significance, likely reflecting limited power with few cohorts. All four approaches performed in a broadly similar range, consistent with immune infiltration being a genuine component of ICB response prediction from bulk transcriptomic data.

To directly quantify the independent predictive contribution of EXPRESSO beyond immune infiltration, we performed a residual analysis in which EXPRESSO scores were regressed against each immune score per cohort and the residuals were used to predict response. Residual EXPRESSO scores retained meaningful predictive ability across all conditions (**Supplementary Table S6**), with residual AUCs ranging from 0.58 to 0.67. In the ICB-non-melanoma cohort set, residual AUCs were consistently higher (0.64–0.67) with lower coefficient of determination (R² = 0.36–0.43) from the linear regression of EXPRESSO score on each immune score, indicating greater independent predictive signal in the pan-cancer setting where immune infiltration alone is less discriminative. Together, these analyses demonstrate that EXPRESSO captures biologically meaningful and clinically relevant transcriptomic signals that extend beyond simple immune cell composition.

### EXPRESSO models generalize beyond training cohorts

To assess whether LOCOCV performance estimates were influenced by overfitting, we performed independent validation for drug categories where additional cohorts could be obtained. Independent cohorts were gathered for five categories: ICB-melanoma (1 cohort), ICB-non-melanoma (3 cohorts including 1 cohort for ICB-lung), trastuzumab (3 cohorts), and chemo-FAC-FEC (15 cohorts). These cohorts were held out entirely from model development. For each drug category, a pooled model was trained on all original cohorts, and each independent cohort was tested against this model. Independent validation ROC-AUCs were concordant with LOCOCV estimates across all five categories (**Figure 4A**), confirming that the models generalize beyond the training data. As a sensitivity analysis, a fold-averaged evaluation approach — in which each independent cohort was tested against every LOCOCV fold model and ROC-AUCs were averaged — yielded comparable results (**Supplementary Figure S9**). For the three drug categories without available independent cohorts (bevacizumab, BRAFi, and paclitaxel), validation relies on LOCOCV performance as reported above.

**Figure 4:**
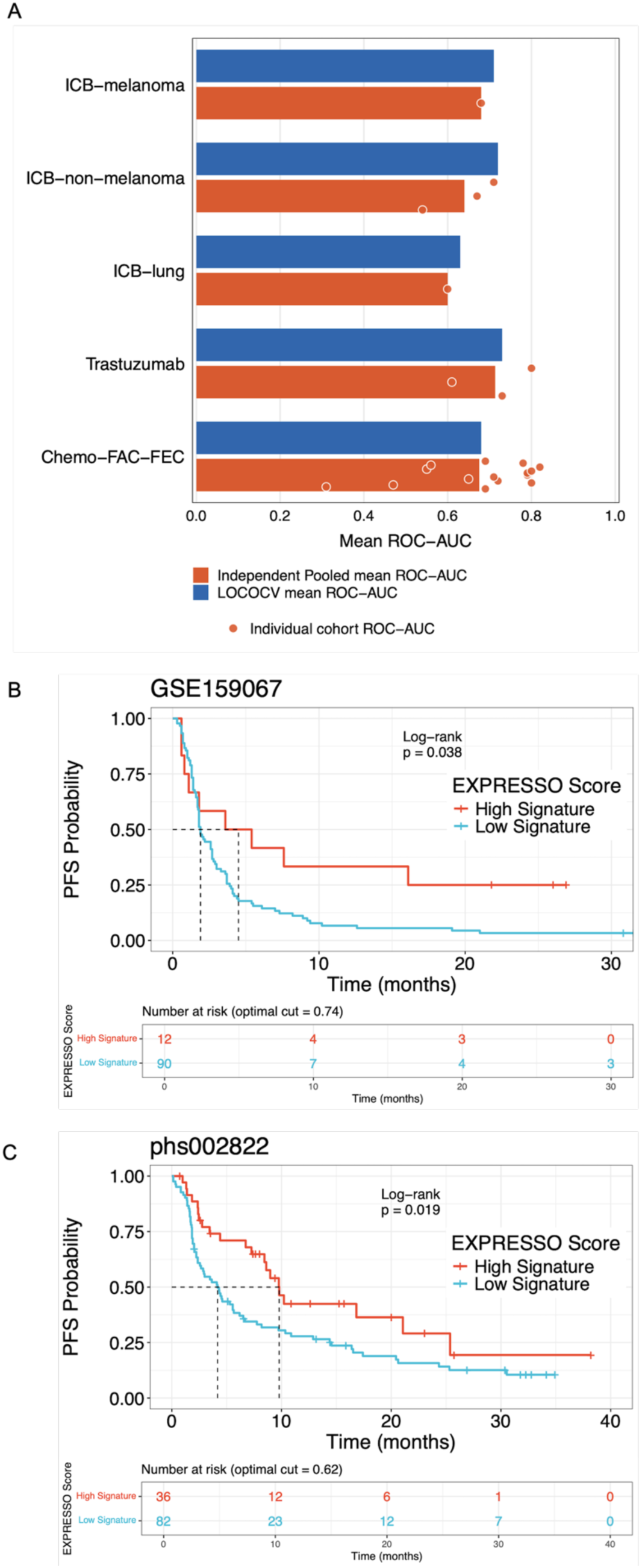
(A) Independent validation of EXPRESSO using held-out cohorts. Horizontal bars show mean ROC-AUC for each drug category under two evaluation strategies: leave-one-cohort-out cross-validation (LOCOCV; blue) and independent validation using a pooled model trained on all original cohorts (orange). Individual cohort ROC-AUCs are overlaid as solid circles with white borders on the independent validation bar. Independent cohorts were available for five drug categories: ICB-melanoma (1 cohort), ICB-non-melanoma (3 cohorts), ICB-lung (1 cohort), trastuzumab (3 cohorts), and chemo-FAC-FEC (15 cohorts). All independent cohorts were held out entirely from model development. ICB-lung is shown as a separate category; the same lung cohort tested against the broader ICB-non-melanoma models yielded lower performance, supporting the rationale for disease-specific modeling (see **Results**). Concordance between LOCOCV and independent pooled validation ROC-AUCs across drug categories indicates that model performance is not driven by overfitting. **(B)& (C) Kaplan-Meier survival curves for EXPRESSO signature score stratification of PFS** in the two largest ICB-non-melanoma cohorts, GSE159067 (B) and phs002822 (C). Patients were stratified into high (red) and low (blue) signature score groups using an optimal cutpoint determined post-hoc by maximizing the log-rank statistic. Numbers at risk are shown below each plot. P-values are from the log-rank test. Median survival lines for each group in each plot are shown by dashed black line.

Among the ICB-non-melanoma independent cohorts, one lung cancer cohort showed lower performance, consistent with the biological heterogeneity of non-melanoma tumor types. This observation motivated the development of a cancer-type-specific model for lung, described below.

To assess whether cancer-type-specific modeling could improve ICB response prediction within the non-melanoma category, we built a specific ICB-lung EXPRESSO model trained exclusively on the five NSCLC cohorts previously included in the broader ICB-non-melanoma model. Prospective validation on a held-out independent NSCLC cohort (GSE274975, n=45) showed improved performance against the ICB-lung model (pooled AUC = 0.60, fold-averaged AUC = 0.64) compared to the ICB-non-melanoma model (pooled AUC = 0.54, fold-averaged AUC = 0.55; **Supplementary Table S7**), supporting the benefit of cancer type-specific modeling. Reactome pathway enrichment of the ICB-lung model’s stable features revealed significant enrichment of complement cascade, inflammasome, and FCGR activation pathways (adjusted p < 0.05), dominated by innate immune activation genes (*FCGR3A, C1QBP, C8B, PYCARD*) — a profile distinct from the adaptive immune and cell cycle progression signatures characterizing the broader ICB-non-melanoma model (**Supplementary Figure S8, Supplementary Table S4**).

The EXPRESSO signature also demonstrated prognostic value beyond binary response classification. In the two largest ICB-treated cohorts with available survival data (GSE159067, head and neck squamous cell carcinoma, n = 102; phs002822, lung cancer, n = 118), the EXPRESSO signature significantly stratified progression-free survival (PFS) (**Figures 4B, 4C**). In GSE159067, the continuous signature score was a significant predictor of PFS by Cox regression (HR = 0.361, Wald p = 0.047), and Kaplan-Meier analysis confirmed significant group separation (log-rank p = 0.038). In phs002822, Kaplan-Meier analysis showed significant PFS stratification (log-rank p = 0.019). The signature did not significantly stratify survival in BRAF/MEK inhibitor cohorts, consistent with the prevalence of universal acquired resistance in this setting. Results from smaller cohorts (n<100) were numerically unstable and are reported in **Supplementary Table S8** for completeness.

### EXPRESSO benchmarks favorably against 20 published gene signatures and transcriptomic predictors

We benchmarked EXPRESSO against a comprehensive set of previously published gene signatures and transcriptomic predictors for treatment response across immune checkpoint blockade (ICB), chemotherapy, and breast cancer therapies. These included ICB-specific signatures such as ADO (49), Myeloid DC (50), TGFB_Mariathasan (51), APM_Thompson (52), APM_Wang (53), B_cell_Budczies and Macrophage (54), CD8_SF (55), CYT (56), Immune Cells (57), IMPRES (58), IPRES (59), COX_IS (60), and TIDE (61); proliferation or oncogenic pathway-related markers such as KDM5A (62) and MKI67 (63); and breast cancer-specific predictors including MammaPrint (64), OncotypeDX (65), Immuno-Oncology signature (66), and an 11-gene signature (67).

Many of these signatures were originally developed in specific therapeutic or disease contexts (e.g., ICB response, chemotherapy sensitivity, or breast cancer prognosis) and were not designed for every drug evaluated here. Nonetheless, we applied the same set of 20 signatures across all six drug classes to establish a uniform benchmarking framework. Per-cohort ROC-AUC values for all 20 published signatures as well as both EXPRESSO variants across all seven drug-specific models are in **Supplementary Table S9**. Across drug-specific models, the best-performing EXPRESSO variant (the mode with the higher mean ROC-AUC) consistently outperformed these existing gene signatures in terms of mean ROC-AUC (**Figures 5A-G**). In ICB-melanoma, the improvement was driven by EXPRESSO-B, whereas EXPRESSO-T did not exceed all the comparators (**Supplementary Figures S10A-E**). For other drugs, both EXPRESSO variants (where available) outperformed published signatures. These results highlight the strength of target-aware supervised modeling with stringent biomarker selection.

**Figure 5:**
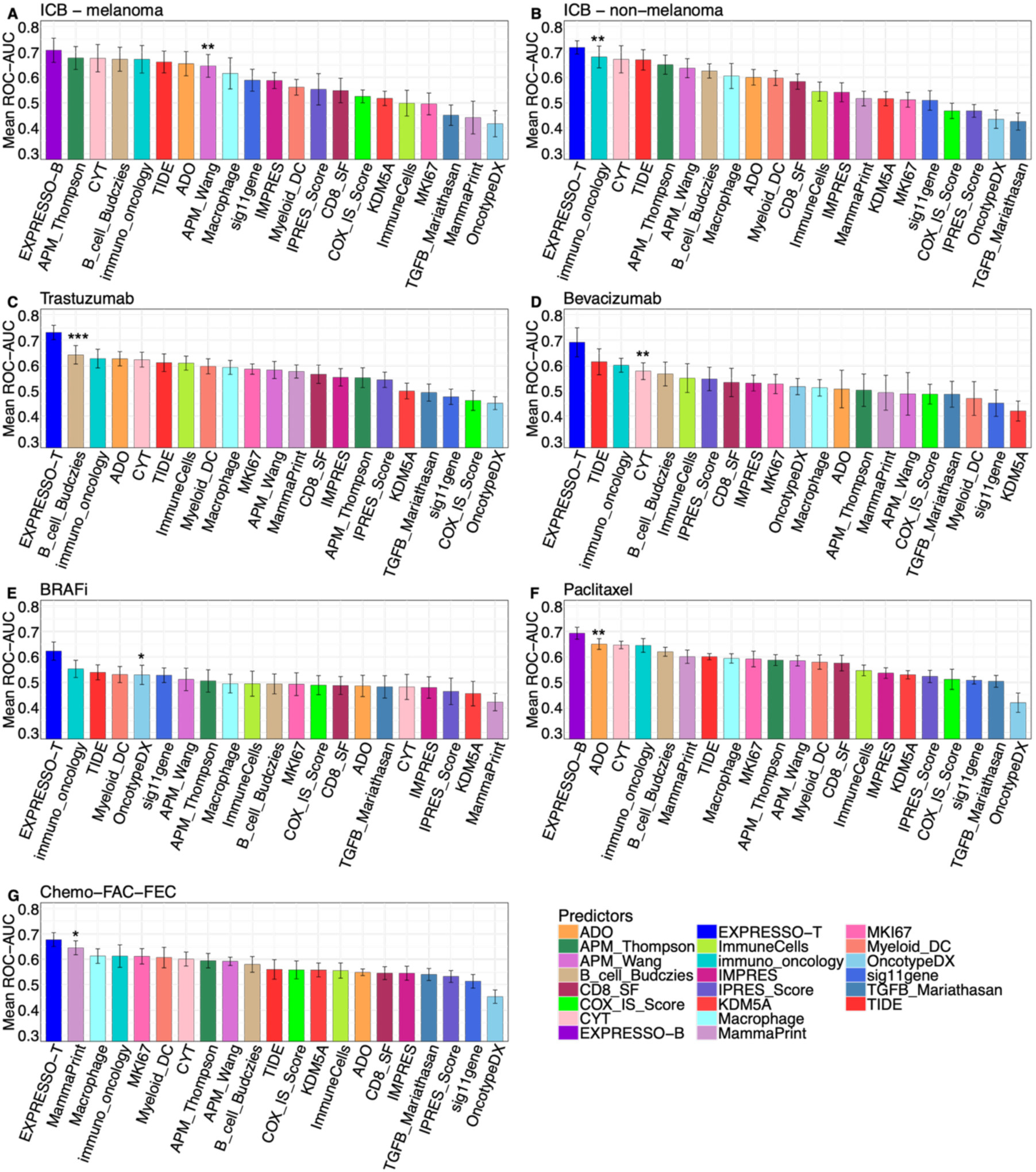
Comparative performance of best-performing EXPRESSO variant for each drug class against existing gene signatures and methods. Bar plots show the mean ROC-AUC values for each predictor across drug-specific models, ordered from highest to lowest mean performance. Significant p-values (one-sided DeLong’s test across all cohorts for a drug) between EXPRESSO variant (by higher mean ROC-AUC between EXPRESSO-T and EXPRESSO-B) and the first significantly different predictor denoted by Asterix (*) with significant levels as follows: * p ≤ 0.1, ** p ≤ 0.05, *** p ≤ 0.01; subsequent predictors with lower performance are also significantly different. While many of the published signatures were originally developed for specific therapeutic or disease contexts, we applied the same set of 20 signatures uniformly across all drugs to enable consistent benchmarking. The best-performing EXPRESSO variant is shown for each drug, as the two EXPRESSO variants achieved comparable performances (bevacizumab and BRAFi have no EXPRESSO-B model), except ICB-melanoma (**Supplementary Figure S10**).

To further validate these findings in fully independent data, we applied the same 20-signature benchmarking framework to the held-out prospective cohorts. For chemo-FAC-FEC—the drug category with the largest number of independent validation cohorts (n = 15) – EXPRESSO maintained the highest mean ROC-AUC among all 20 published signatures and methods evaluated on the same held-out data (**Supplementary Figure S11**). For the remaining drug categories where independent cohort numbers were small (1–3 cohorts), a reliable multi-method comparison was not feasible. Together, these results underscore the importance of treatment-specific, target-aware predictors in precision oncology.

### Supervised prediction capability limit of transcriptomic data varies across different treatment types

To estimate the upper bound of supervised predictive performance and assess the impact of training data size, we conducted incremental training experiments (**Figures 6A-G**). For each drug category, we sequentially added training cohorts one at a time (excluding the test cohort) in various orders and retrained the best performing EXPRESSO variant (the mode with the higher mean ROC-AUC) at each step of each order (**Methods**). Across drug types, we observed consistent increases in ROC-AUC as additional cohorts were incorporated, although in some drug types, the improvements diminished after a certain point—suggesting saturation of predictive performance for some drugs.

**Figure 6:**
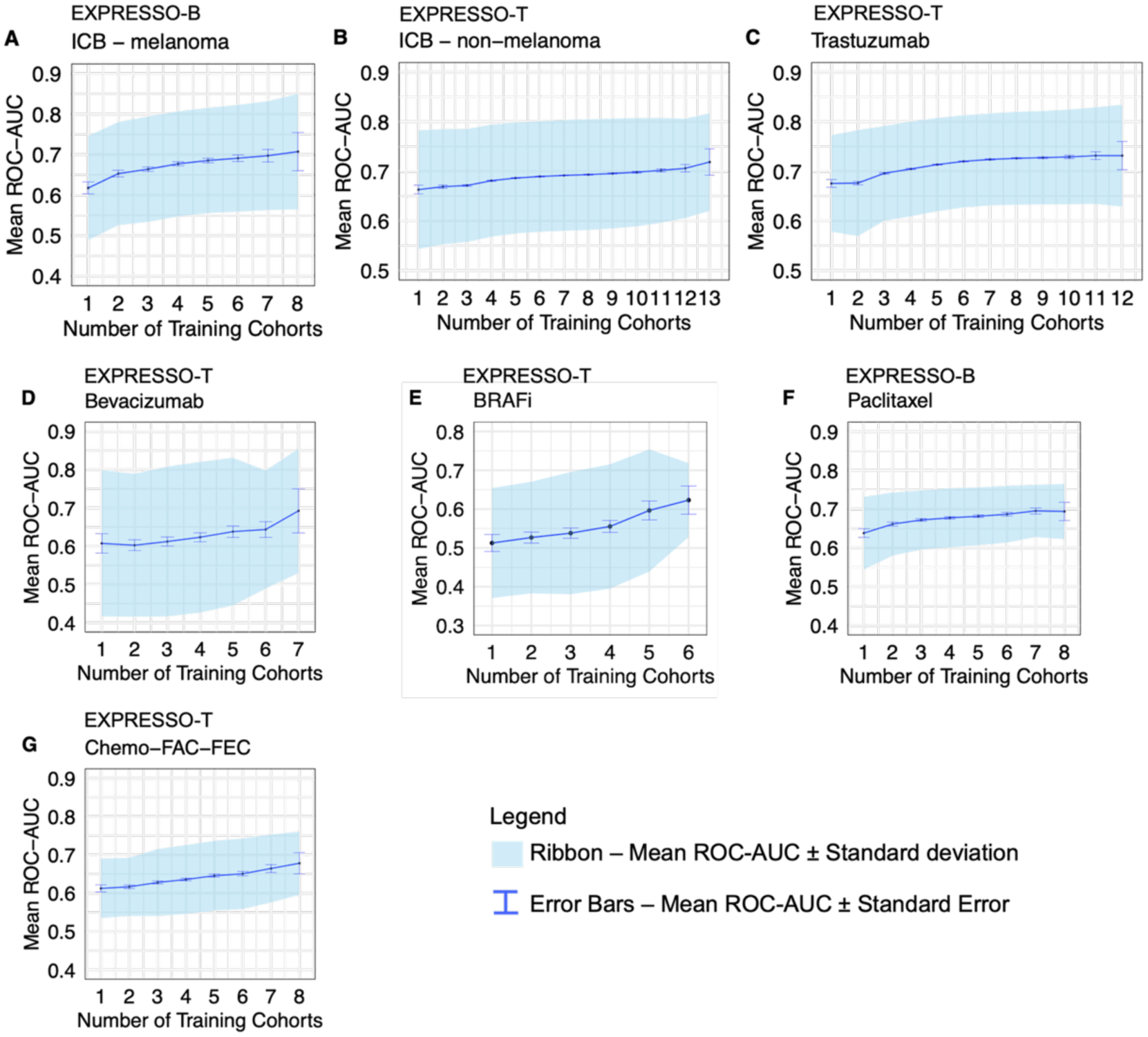
Line plot with standard deviation as ribbon (sky blue) and mean standard error as error bars for the upper bound of transcriptome-based prediction for seven drug-specific best-performing EXPRESSO variants (either EXPRESSO-T or EXPRRESSO-B, as noted on top of the plots) for each drug class.

In the analyses shown in **Figure 6**, each training subset with the same number *t* of training cohorts was treated equivalently, but these subsets may have rather different sample sizes. Therefore, we also made scatter plots of sample size (x-axis) against ROC-AUC (y-axis) for fixed numbers of training cohorts (**Methods, Supplementary Figures S3A-G**). Reassuringly, ROC-AUC is consistently positively correlated with sample size, significantly for some drugs and insignificantly for other drugs. The lines of best fit have small coefficients for x suggesting that where EXPRESSO prediction accuracy has not plateaued, the addition of new cohorts of roughly average size (relative to the current cohorts) would help.

In summary, these experiments suggest that predictive performance may have reached its upper bound for trastuzumab, ICB-melanoma, and ICB-non-melanoma where we have sufficient (at least nine) cohorts. In contrast, models for bevacizumab, BRAF inhibitors, and for chemotherapy cohorts have not yet plateaued.

## DISCUSSION

In this study, we evaluated the predictive potential of bulk transcriptomic data across diverse cancer therapies using supervised machine learning frameworks. By compiling and uniformly processing 69 clinical cohorts (3,729 patients) covering six frontline drug classes – anti-PD-1 immunotherapy (melanoma and non-melanoma), trastuzumab, bevacizumab, BRAF inhibitors, paclitaxel, and FAC/FEC chemotherapy – we developed an approach to assess transcriptomics in a pan-cancer setting. Notably, all datasets are publicly available (but some requiring dbGaP permission) and the Expresso code is available on Zenodo.

We showed that expression of drug targets can predict treatment response quite reasonable (mean ROC-AUC of 0.68-0.69) for selected monoclonal antibody therapies (trastuzumab, bevacizumab, and anti-PD-1 inhibitors for non-melanoma cancers), but not for small-molecule BRAF inhibitors, paclitaxel, or FAC/FEC chemotherapy. In melanoma, PD-L1 expression was not a definitive predictor of response (68), though we observed modest predictive value (mean ROC-AUC 0.65). These observations suggest that monoclonal antibody efficacy may depend more directly on target abundance (45), whereas chemotherapy and small-molecule inhibitors often act through mutated proteins or multiple/off-target mechanisms, making single-gene transcript levels an imperfect proxy for drug activity (69). While previous studies have reported target-specific associations in narrower drug–cancer-cohort contexts(14,70), our work provides a broader evaluation across multiple cancer types, therapies, and clinical cohorts.

We developed EXPRESSO, a drug-specific LASSO framework, integrating biological priors with whole transcriptome features to capture predictive signals beyond target expression while reducing risk of overfitting using LOCOCV strategy. It incorporates biological information in two variants: EXPRESSO-T, which includes only the drug’s target gene(s), and EXPRESSO-B, which additionally includes context-specific biomarkers, as penalty-free features for both. Both variants performed comparably for all the drugs, except in ICB-melanoma, where EXPRESSO-B outperformed EXPRESSO-T (p=0.02, one-sided DeLong’s test). While achieving higher performance than 20 published gene signatures and prediction tools, overall, EXPRESSO achieved moderate response prediction across several drug types. In many cases, it consistently outperformed the baseline supervised vanilla model without biological priors, demonstrating the value of incorporating biological knowledge. Benchmarking against seven alternative machine learning methods under identical LOCOCV conditions further confirmed that EXPRESSO’s regularized linear framework is well-suited to this prediction problem, matching or exceeding nonlinear methods in five of seven drugs without sacrificing interpretability.

Independent prospective validation across five drug categories confirmed that LOCOCV performance estimates were not driven by overfitting. Independent validation ROC-AUCs were concordant with LOCOCV estimates, and the cancer-type-specific ICB-lung model showed improved performance over the broader ICB-non-melanoma model on a held-out NSCLC cohort, supporting the value of disease-specific modeling. For chemo-FAC-FEC, where 15 independent cohorts were available, EXPRESSO also maintained the highest mean ROC-AUC among 20 published signatures on held-out data, further corroborating the robustness of its predictions beyond the cross-validation framework. The survival analyses provide preliminary evidence that the EXPRESSO signature has prognostic value in ICB-non-melanoma cohorts beyond binary response prediction, particularly in larger cohorts.

This study has several limitations. First, using both RNA-seq and microarray datasets may introduce technical variability. We applied batch correction but observed no improvement in predictive performance; rank normalization was therefore used as the sole harmonization step. We showed in **Supplementary Results S2** and **Supplementary Figures 2C-D** that this choice does not leave a significant residual platform bias between microarrays vs. RNA-seq. Second, anti-PD-1 and anti-PD-L1 therapies were analyzed together to increase cohort sizes, despite their known efficacy differences(71). Third, importantly, many therapies are often delivered in combinations, but we modeled only the primary treatment to maximize cohort inclusion. Fourth, the Kaplan–Meier analyses shown in **Figures 4B and 4C** used a cutpoint selected post hoc to maximize the log-rank statistic, rather than a pre-specified threshold such as the median signature score. Because the cutpoint is chosen from the same data used for testing, the resulting log-rank p-values are optimistically biased, and these analyses should be regarded as exploratory. The Cox proportional hazards analyses, which use the continuous signature score and require no dichotomization, are not subject to this bias. Prospective confirmation in independent cohorts using a pre-specified cutpoint will be required before the signature could be applied for survival stratification. Fifth, one may ask whether expression-based ICB response goes beyond just reflecting association with immune infiltration. While EXPRESSO’s top selected genes include established markers of T cell infiltration and IFN-γ signaling, CIBERSORT deconvolution revealed that EXPRESSO scores correlate most strongly with M1 macrophage polarization rather than CD8^+^ T-cell infiltration alone, suggesting the model captures a broader immune activation signature. Notably, EXPRESSO outperformed TIS, *CD8A* expression, and ESTIMATE immune score in direct AUC comparison, and residual analysis confirmed that EXPRESSO retains independent predictive value beyond what these simpler immune scores capture, particularly in the pan-cancer setting. A further limitation concerns the independent verifiability of the EXPRESSO-B biomarker selection procedure. Because biomarkers are identified within each outer LOCOCV fold using nested cross-validation strictly confined to training data, the selected gene sets are necessarily fold-specific — an intended feature that can appear counterintuitive (see **Supplementary Materials S2**). We have deposited the complete biomarker selection pipeline to our Zenodo repository **(Data/Code availability section)** to enable independent verification. Finally, we posed prediction dichotomously (“Will patient *p* respond to treatment *t*?”), whereas tumor boards face a combinatorial decision across multiple treatment options.

Perhaps the most important finding of this study is a negative one: at least in some treatment types, prediction accuracy plateaued and did not improve with additional training cohorts. This suggests that supervised bulk transcriptomics alone may be insufficient for robustly predicting patients’ response in highly heterogeneous cohorts, especially for the case of non-targeted therapies. Our findings highlight the inherent limitations of supervised brute-force learning for certain treatments, while underscoring the potential of incorporating additional data and deeper mechanistic modeling of tumor heterogeneity to improve transcriptomics-based predictors. Future directions are of two conceptual dimensions: (a) developing bulk expression models for still larger and more clinically homogenous datasets, that, importantly, better model and incorporate the underlying biological mechanisms of treatment resistance and response, and (b) bringing additional omics data to the mix, include integrating DNA mutation data, prior treatments information, clinical metadata, and richer multi-omics (e.g., single-cell transcriptomics, proteomics, spatial immune profiling and others), to provide a more comprehensive view of therapeutic response mechanisms. Increasing access to large, well-annotated, and diverse clinical trial cohorts, with such deeper molecular profiling, will hence be important for advancing data-driven precision oncology. Fortunately, recent developments in AI-based inference of such types of data in a fast and cost-effective manner directly from the tumor pathology slides (e.g., DeepPT (72), Path2Omics (73), Path2Space (74) may facilitate the latter in a readily accessible and cost effective manner.

The Hallmarks of Predictive Oncology (75) offer a useful framework for situating EXPRESSO within the broader landscape of clinical prediction models. Regarding **Data Relevance and Actionability**, EXPRESSO was built and validated exclusively on pre-treatment transcriptomic data from clinical trial cohorts, addressing a directly actionable clinical question. On **Expressive Architecture**, we intentionally favored a parsimonious, biologically constrained LASSO model over more complex architectures, prioritizing robustness and generalizability given the modest sample sizes of individual cohorts. **Standardized Benchmarking** is a core strength: EXPRESSO was evaluated via rigorous LOCOCV across 69 cohorts and benchmarked against 20 published signatures and baseline models. **Generalizability** is both a strength and a recognized limitation — performance was consistent across cancer types for targeted therapies, but saturation analyses revealed that for non-targeted chemotherapies and heterogeneous drug classes, predictive accuracy plateaued regardless of additional training data, indicating fundamental constraints of bulk transcriptomics alone. **Interpretability** is well-served by the sparse LASSO framework, whose features — drug target genes and a small number of biologically motivated biomarkers — are directly mechanistically interpretable. **Accessibility and Reproducibility** are fully addressed: all datasets are publicly available (some via dbGaP), and the complete EXPRESSO codebase is deposited on Zenodo **(Data/Code availability section)**. Finally, regarding **Fairness**, we acknowledge that the cohorts analyzed were largely drawn from clinical trials that may not fully reflect the demographic and geographic diversity of real-world cancer patients, a limitation that future studies with broader representation should address.

## ACKNOWLEDGEMENTS

This research was supported in part by the Intramural Research Program of the National Institutes of Health (NIH), National Cancer Institute. The contributions of the NIH author(s) were made as part of their official duties as NIH federal employees, are in compliance with agency policy requirements, and are considered Works of the United States Government. However, the findings and conclusions presented in this paper are those of the author(s) and do not necessarily reflect the views of the NIH or the U.S. Department of Health and Human Services. This work used the computational resources of the NIH HPC Biowulf cluster (http://hpc.nih.gov). We thank Stacy Brody and Alicia Livinski from the NIH Library and Gal Dinstag from Pangea, for their help with this work. The authors used ChatGPT 4o and 5 to improve the quality of writing of this manuscript.

## AUTHOR CONTRIBUTIONS

**L.R.P.:** Data collection, data pre-processing, data curation, conceptualization, software, methodology, formal analysis, visualization, writing the draft. **E.M.G.:** Software, methodology, formal analysis, visualization. **N.U.N:** Conceptualization, methodology, writing the draft. **S.M.:** Data pre-processing, formal analysis. **S.P.:** Data collection, Formal analysis. **T.C.:** Data pre-processing, software. **E.M.C.:** Data pre-processing. **T-G.C:** Data pre-processing. **S.R.D.:** Data collection. **Y. K.:** Data collection. **E.D.S.:** Methodology. **P.S.R.:** Data collection. **D-T.H.:** Methodology review. **S.H.:** Methodology review and carried out the revisions. **A.A.S.:** Data collection, data acquisition, supervision, edited the original draft and carried out the revisions. **E.R.:** Conceptualization, data collection, supervision, project administration, funding acquisition, methodology, edited the original draft and carried out the revisions. All authors critically reviewed and approved the final version of the manuscript.

## Supplementary Materials and Supplementary Figures for EXPRESSO paper

### Supplementary Materials S1

#### Generating LASSO Logistic Regression Models

We utilized the glmnet R package (1) to create LASSO logistic regression models from the rank-normalized features. Samples from each set of training cohorts were pooled to form a single training dataset, and the optimal shrinkage parameter was determined using 10-fold cross-validation (default option) via the cv.glmnet routine. We set the family parameter to “binomial” to generate logistic models. To balance the sensitivity and specificity of predictors, sample weights were adjusted so that responders and non-responders contributed equally to the total weight. Besides these settings, we used default parameters of cv.glmnet.

#### Leave-One-Cohort-Out Cross-Validation (LOCOCV) Framework

To ensure robust and generalizable evaluation, we employed a Leave-One-Cohort-Out Cross-Validation (LOCOCV) framework for the supervised learning. In this setup, the number of folds equals the number of available cohorts, with each fold holding out one cohort as an independent test set while training on the remaining cohorts. This approach directly models real-world clinical scenarios, where predictive models are applied to unseen patient populations from distinct clinical settings.

#### EXPRESSO – T (Target-only) model

EXPRESSO is a drug-specific machine-learning-based supervised learning framework specifically tailored for transcriptome-based drug response prediction. EXPRESSO leverages LASSO regression with L1 regularization, chosen for its capability to perform automatic feature selection and mitigate overfitting.

EXPRESSO incorporates the entire transcriptome as input for feature selection and model training. EXPRESSO-T models were constrained to include the relevant target genes (as regularization free) for each drug: *ERBB2* for trastuzumab, *VEGFA* for bevacizumab, *BRAF* for BRAFi, *CD274* (PD-L1) for anti-PD-1 checkpoint inhibitors for melanoma and other non-melanoma, *TUBB1* for Paclitaxel and *TOP2A* and *TYMS* for Chemo-FAC-FEC (cyclophosphamide, doxorubicin, fluorouracil / cyclophosphamide, epirubicin, fluorouracil). For anti-PD1 therapy, we chose *CD274 (PD-L1)* as the target gene as anti-PD1 therapy targets the interaction between PD-1 and PD-L1, and PD-L1 is the ligand which is expressed in cancer cells.

The EXPRESSO framework applies the Leave-One-Cohort-Out Cross-Validation (LOCOCV) setup, with a training set consisting of gene expression data, drug response labels, and the identified target gene for a specific drug; the last of these design choices is what distinguishes EXPRESSO-T. It uses all background genes for normalization and uses the normalized expression values to train a LASSO logistic regression model. LASSO chooses a small set of genes as features from the input. We used the 0.5 threshold to differentiate predicted responders from non-responders. For the independent test sample, EXPRESSO-T applies the same normalization rules, the trained logistic regression model, and cutoff to predict response.

#### EXPRESSO-B adds biomarkers from Differential Expression Analysis

In the EXPESSO-B mode, biomarkers were identified independently within each fold of the outer LOCOCV loop to avoid information leakage from the test cohort. In each outer fold, one cohort was held out as the test set, while the remaining cohorts formed the training set. To robustly identify biomarkers from the training data alone, we implemented a nested LOCOCV within these training cohorts; each training cohort is logically associated with one iteration of the inner loop. Each inner fold held out one cohort as a validation set, while the remaining cohorts were used for differential expression (DE) analysis. Within each inner training fold, gene expression data from responders and non-responders were compared using the ‘limma’ package with default settings (2) on normalized expression data.

From each inner fold, we extracted p-values and average log fold changes for all genes. DE p-values from inner CV folds were combined using an empirical adaptation of Brown’s method (3). For each drug category, the covariance among cross-validation folds was estimated once, genome-wide, from the matrix of per-fold log2 fold-changes across all candidate genes — rather than per gene — capturing the correlation induced by overlapping training cohorts across the nested LOCOCV folds. This shared fold-by-fold covariance matrix was then used to combine each gene’s own vector of per-fold DE p-values via a moment-matching (effective degrees-of-freedom) approach, rather than treating folds as independent as under Fisher’s method. Genes were ranked by the combined p-values, and a Benjamini–Hochberg false discovery rate (FDR) correction was applied. Genes with FDR < 0.05 and mean absolute log fold change > 0.4 were considered for further pruning.

To further refine the candidate list of differentially expressed genes, we evaluated the predictive utility of each candidate gene in ascending order by FDR-adjusted p-value. For each gene, we computed the change in ROC-AUC (ΔAUC) when it was added (along with the target gene) as a regularization-free feature in the LASSO model, compared to the baseline target-only model.

Genes were retained only if they improved ROC-AUC by at least 0.02 and had a significant Wilcoxon rank-sum p-value comparing ROC-AUCs across folds (target-only, EXPRESSO-T vs. target+biomarker, EXPRESSO-B model). From these, at most the three top-ranking genes per fold that met both criteria were selected for inclusion in the model; if fewer than three genes qualified, all available candidates were included. If no candidate genes passed both ΔAUC and significance criteria in a fold, the model defaulted to the target-only configuration.

The final list of selected genes—specific to each drug and outer fold—was passed into the LASSO model along with the drug target as regularization-free features. Importantly, all biomarker discovery steps were confined to training data only, maintaining strict separation from the test cohort.

#### More details on the EXPRESSO-B (Target + Biomarker) model

In the EXPRESSO-T model, gene expression values for all background genes were provided as candidate predictors, but only the known drug target gene was designated as *regularization-free* (i.e., exempt from penalization in the LASSO). This setup establishes a biologically grounded baseline by allowing the model to prioritize the drug’s direct target while applying penalization to the rest of the transcriptome.

In contrast, in the EXPRESSO-B (target + biomarker) model, we retained the drug target gene as regularization-free but additionally included a set of cohort-specific biomarkers identified as described in the preceding subsection as unpenalized features. The remaining genes were subject to regularization as in EXPRESSO-T. Both models were implemented under the same Leave-One-Cohort-Out Cross-Validation (LOCOCV) setup.

#### Two types of baseline models

As one baseline, we implemented a supervised model in which the entire transcriptome served as input features for each drug using a LASSO framework. Since different cohorts profiled partially overlapping gene sets, we restricted the analysis to genes measured in at least three cohorts to enable consistent cross-cohort comparisons.

For a second, biologically informed baseline, we assessed whether the expression of each drug’s known target gene alone could predict patient response in an unsupervised, model-free manner. For each drug and cohort, patients were ranked by their pre-treatment normalized expression of the corresponding target gene. We then evaluated prediction performance by calculating the area under the receiver operating characteristic curve (ROC-AUC). This analysis served as a reference point for assessing the added value of supervised modeling compared to unsupervised modeling.

#### Performance measures

To evaluate model performance in each fold of the Leave-One-Cohort-Out Cross-Validation (LOCOCV) framework, we used both discrimination and effect-size–based metrics. The primary evaluation metric was the area under the receiver operating characteristic curve (ROC-AUC), which quantifies the model’s ability to distinguish responders from non-responders across varying decision thresholds. ROC-AUC was computed using the predicted probability scores from the model on the held-out test cohort in each fold.

To complement the ROC-AUC, we also calculated the odds ratio (OR) to quantify the enrichment of predicted responders among true responders in the test cohort. Patients were ranked by their predicted scores, and a threshold of 0.5 was used to define the predicted-positive group. The odds of true response in this group versus the rest of the test cohort were used to compute the OR, along with a 95% confidence interval using the Fisher’s exact test.

In addition to per-fold metrics, we computed the mean ROC-AUC and median OR across all folds for each drug to assess the overall predictive performance and compare between the models. The median value of OR across cohorts for each drug class was reported to minimize the impact of outliers. For statistical comparison of model performance, we used paired Wilcoxon signed-rank tests across folds and drugs and one-sided DeLong’s tests to compare ROC-AUCs of nested models, where applicable.

#### EXPRESSO models benchmarking against other signatures and predictors

For benchmarking against the 20 published gene signatures and transcriptomic predictors, signature scores were computed using the same rank-normalized expression data used throughout the study (rank normalization across genes per patient). For signatures requiring a dedicated computational framework, scores were computed as follows: TIDE(4) scores were computed using the published TIDE framework, which estimates T cell dysfunction scores in tumors with high cytotoxic T lymphocyte (CTL) infiltration and T cell exclusion scores in tumors with low CTL levels, integrating these into a final TIDE score predictive of ICB response. IMPRES(5) scores were calculated based on 15 predefined pairwise gene expression relationships between immune checkpoint genes, with the final score defined as the total number of gene pairs satisfying the specified directional expression criteria. IPRES(6) scores were computed as the mean expression of genes included in the published innate anti-PD-1 resistance signature, following appropriate normalization to ensure comparability across genes and samples. COX-IS(7) scores were calculated by quantifying the relative expression of predefined cancer-promoting (CP) and cancer-inhibitory (CI) gene sets, reflecting the balance between opposing inflammatory programs in the tumor microenvironment. For all remaining multi-gene signatures (e.g., ADO, CYT, APM_Thompson, MammaPrint, OncotypeDX), scores were computed as the mean of the rank-normalized expression values of the constituent genes available in each cohort. For single-gene predictors (e.g., *MKI67, KDM5A*), the rank-normalized expression of that gene was used directly as the score. ROC-AUC was computed per cohort by ranking patients by their signature score and evaluating discrimination against the binary response label, consistent with the evaluation approach applied to EXPRESSO throughout.

#### EXPRESSO models benchmarking against other machine learning models

To evaluate the contribution of nonlinear relationships to drug response prediction, EXPRESSO was benchmarked against seven alternative machine learning methods: Random Forest (500 trees), XGBoost (300 estimators, learning rate 0.05), SVM with RBF and linear kernels, KNN (k=5), shallow Neural Network (128-64 hidden units), and a deep MLP (256-128-64 hidden units, PyTorch). All methods used identical input features, normalization, and LOCOCV framework as EXPRESSO. Class imbalance was addressed using balanced sample weights for all applicable methods. Statistical comparison used the two-sided Wilcoxon signed-rank test with EXPRESSO as reference, and 95% confidence intervals were estimated by bootstrap resampling (2,000 iterations). Comparisons with other machine learning methods were implemented in Python 3.11 using scikit-learn(8) (v1.6). XGBoost was implemented using the xgboost Python package(9) (v2.1.4). The deep MLP was implemented in PyTorch(10) (v2.5.1+cu121) with batch normalization, dropout (rate=0.3), and Adam optimizer(11).

#### CIBERSORT deconvolution and immune score benchmarking

CIBERSORT(12) deconvolution was performed on all ICB cohort expression matrices using the LM22 signature matrix with 100 permutations and quantile normalization enabled. Expression matrices were provided in non-log TPM format as required by CIBERSORT. Estimated cell-type fractions were merged with sample-level EXPRESSO prediction scores generated by leave-one-cohort-out cross-validation, and Spearman correlations were computed between EXPRESSO scores and each of the 22 LM22 cell-type fractions. P-values were adjusted for multiple comparisons using the Benjamini-Hochberg false discovery rate procedure(13).

For benchmarking, three simpler immune scores were computed per sample using the same ranked normalization pipeline as EXPRESSO. The Tumor Inflammation Signature (TIS) score(14) was computed as the mean of available TIS genes (18-gene panel) across samples after ranked normalization. The ESTIMATE immune score was computed as the mean of available genes from the Immune141_UP gene set(15). *CD8A* expression was extracted as a single-gene score from the ranked-normalized matrix. Per-cohort AUC (responder vs non-responder) was computed for each score using the pROC package in R.

To assess whether EXPRESSO provides predictive value independent of immune infiltration, a residual analysis was performed. For each cohort, EXPRESSO scores were regressed against each immune score using linear regression, and the residual scores were used to recompute AUC. The coefficient of determination (R²) from each regression quantifies the proportion of EXPRESSO score variance explained by the respective immune score.

#### Incremental Training for Upper-Bound Performance Estimation

To assess the upper bound of prediction capability and the impact of training cohort size on model performance, we conducted an incremental training analysis within the LOCOCV framework. For each test cohort, we iteratively increased the number of training cohorts by one at a time—from a single cohort up to all available training cohorts excluding the test set—and trained the model at each step. At each increment, the model was retrained from scratch using the expanded training set and evaluated on the fixed test cohort. The experiments were done with many orders of adding the training cohorts one at a time.

This stepwise accumulation allowed us to quantify how prediction performance (measured by ROC-AUC) evolves as more training data becomes available. By plotting ROC-AUC as a function of number of training sets for each drug, we estimated the trajectory and potential saturation point of predictive performance.

In the main experiment, different runs with *t* training cohorts for each value of *t* = 1, 2, 3,…were treated as interchangeable. For those settings in which performance may not have plateaued, it may be useful to estimate how many additional patient samples (not data sets) would get the modeling to a performance plateau. Therefore, we plotted the prediction performance (ROC-AUC, y-axis) as a function of the number of patients in the training subset (x-axis) in Supplementary Figure S3A-G.

#### Gene Annotation Sources

For RNA-seq quantification using Salmon(16) (v1.11.4, default parameters), transcript-level abundances were estimated against the [GENCODE v43 / Ensembl release109] human reference transcriptome (GRCh38). Gene symbols were used as the common identifier across platforms; genes with ambiguous or deprecated symbols were excluded from cross-cohort analyses.

#### Platform Harmonization (Microarray vs. RNA-seq)

To harmonize expression data across microarray and RNA-seq platforms, expression values from each platform were processed separately before rank normalization. For microarray cohorts, raw or pre-normalized intensity values from GEO were used as provided by the submitting laboratory. Probes with zero expression values were first removed, and any remaining negative expression values were set to zero. For RNA-seq cohorts, TPM files were downloaded from GEO as-is; Ensembl or Entrez gene IDs were converted to NCBI gene symbols using the biomaRt R package(17) (v2.66), and only genes with a successful ID mapping were retained.

Both data types were then subjected to within-sample rank normalization – which places all samples on a common ordinal scale regardless of the original measurement technology, effectively removing platform-specific distributional differences without requiring explicit batch correction. We also evaluated frozen surrogate variable analysis(18) (fsva) for batch correction but found it did not improve prediction performance. (see limitations paragraph under Discussion).

#### Filtering Criteria and Gene Counts

Prior to normalization, expression data were processed separately by platform. For RNA-seq cohorts, TPM files were downloaded from GEO. For microarray cohorts, probes with zero expression values were removed. Negative expression values, which can arise in microarray data from background subtraction steps during array preprocessing, were set to zero as they do not represent meaningful biological signal. The sets of genes analyzed varied across drug classes, depending on the training cohorts available for each model. During model training, only genes present in at least three cohorts within a given drug class were considered, ensuring a minimum level of cross-cohort representation. Genes not meeting this threshold were excluded from the background gene set and were not used in model building.

#### Rationale for Rank Normalization

Rank normalization was chosen as the primary cross-platform harmonization strategy for several reasons. First, it is a non-parametric transformation that preserves relative expression ordering within each sample without making distributional assumptions — an important property when integrating microarray intensities and RNA-seq TPM values, which differ fundamentally in scale and dynamic range. Second, because normalization is applied within each patient independently (across genes), it does not require paired samples or a reference distribution, making it applicable to cohorts of varying size. Third, and empirically, we evaluated standard batch correction approaches (e.g., fsva(18)) and found they did not improve downstream predictive performance (see Limitations), consistent with prior reports that rank-based methods are robust across transcriptomic platforms(19).

#### Software Versions, Key Packages, and Parameters

All analyses were performed in R (v4.5.2). Key packages included: biomaRt(17) (v2.66), WGCNA(20) (v1.74, collapseRows with method = “MaxMean”), tximport(21) (v1.38 default settings), and Salmon(16) (v1.11.4, default parameters). Rank normalization was performed using base R’s rank() function with ties.method = “average”, with ranks subsequently scaled by the total number of genes.

Model training and regularized feature selection were performed using the glmnet R package(22), implementing a LASSO penalized logistic regression model (family = “binomial”, alpha = 1 by default in cv.glmnet). The optimal regularization parameter lambda was selected using 10-fold cross-validation (nfolds = 10), with the minimum cross-validation error criterion (lambda.min). To handle the class imbalance between responders and non-responders, inverse-proportion response weights were applied during model training. In the EXPRESSO-T variant, the drug target gene(s) and in the EXPRESSO-B variant, identified biomarkers together with the drug target gene(s) were included as unpenalized features (penalty.factor = 0) to ensure their retention in the model regardless of regularization. For EXPRESSO-B, the unpenalized feature set in each outer fold comprised up to three biomarker genes selected by the ΔAUC criterion (max_biom = 3) together with the drug target gene(s) for the relevant drug class. Prior to model fitting, the expression matrix was standardized using base R’s scale() function (zero mean, unit variance). All model training used a fixed random seed for reproducibility.

Comparisons with other machine learning methods were implemented in Python 3.11 using scikit-learn(8) (v1.6). XGBoost was implemented using the xgboost Python package(9) (v2.1.4). The deep MLP was implemented in PyTorch(10) (v2.5.1+cu121) with batch normalization, dropout (rate=0.3), and Adam optimizer(11).

### Supplementary Materials S2

#### AUC sensitivity analysis for cohort size

To assess whether the ≥10 patient threshold introduces instability or bias into performance estimates within the LOCOCV framework, we conducted a sensitivity analysis varying the minimum cohort size threshold from 10 to 30 patients (in steps of 5) across all applicable drug cohorts. This sensitivity analysis was applied to ICB-melanoma, ICB-non-melanoma, trastuzumab, bevacizumab, and BRAFi, as the paclitaxel and chemo-FAC/FEC datasets contain at least 30 patients (**Supplementary Figures S2A-B**).

For each threshold, mean area under the curve (AUC) and median odds ratios (OR) were computed across test cohorts whose size was at least that threshold. Confidence intervals (95%) were obtained by bootstrap resampling performed at the cohort level, where cohort AUC values were resampled with replacement (2,000 iterations), quantifying uncertainty in the all-cohort mean AUC arising from between-cohort variability. For bevacizumab and BRAFi, which have fewer cohorts overall, reliable bootstrap estimation was feasible only up to the threshold at which at least 5 cohorts remained; higher thresholds were excluded from the plot as the resulting estimates become unreliable due to insufficient cohort count rather than true model instability.

Point estimates of mean AUC remained stable across all thresholds for all five drug cohorts. For trastuzumab, mean AUC varied by less than 0.01 across thresholds from ≥10 to ≥30 patients (n=13 to n=8 cohorts retained), with fully overlapping confidence intervals throughout — representing the strongest evidence of robustness (**Supplementary Figure S2A-B**). For ICB-melanoma and BRAFi, mean AUC varied by ≤0.04 with overlapping CIs across all thresholds. For ICB-non-melanoma and bevacizumab, mean AUC varied by ≤0.06, with the larger variation occurring only at the highest thresholds (≥25, ≥30) where as few as 5 cohorts remained; at these thresholds, 95% CIs continued to overlap with lower-threshold estimates, indicating that the observed shift in mean AUC at these thresholds is a consequence of averaging over fewer cohorts. Median OR point estimates remained consistently above 1.0 across all thresholds and all drug cohorts, supporting directional discrimination at every threshold tested; the wider median OR confidence intervals at higher thresholds are consistent with the known sensitivity of Fisher’s exact test to small cell counts as cohort numbers decrease. In all cases, 95% confidence intervals overlapped across thresholds, indicating no statistically distinguishable difference in mean AUC performance. The ≥10 threshold was selected to maximize the number of cohorts available for LOCOCV training set construction, rather than being informed by predictive performance.

Taken together, these results demonstrate that the ≥10 patient threshold does not materially affect our conclusions, and that EXPRESSO’s performance estimates are not driven by the inclusion of small test cohorts in the LOCOCV framework.

#### AUC sensitivity analysis for sequencing platform

Because our cohort collection combines microarray– and RNA-seq-profiled samples, harmonized only by within-patient rank normalization, we assessed whether residual platform-specific technical variation systematically influences per-cohort model performance. Across all 91 cohorts (39 RNA-seq, 52 microarray), mean ROC-AUC was 0.68 for RNA-seq and 0.71 for microarray cohorts; this difference was not statistically significant (Wilcoxon rank-sum test, p=0.50) (**Supplementary Figure S2C**). Restricting to the 69 LOCOCV cohorts, the result was unchanged (RNA-seq mean=0.69, microarray mean=0.71; p=0.57).

Because platform usage varies by drug class, a linear model of AUC as a function of platform, adjusting for drug class (AUC ∼ Platform + Drug), was additionally fit. The estimated coefficient for platform effect (microarray relative to RNA-seq) was 0.039 (p = 0.19) and remained non-significant when the model was weighted by cohort size (platform effect coefficient = 0.048, p = 0.14) (Supplementary Figure S2D). Notably, the direction of the platform difference was not consistent across drug classes — RNA-seq cohorts scored higher for trastuzumab and ICB-non-melanoma, while microarray cohorts scored higher for the remaining drug classes — consistent with residual noise rather than a systematic platform artifact.

Finally, to test whether any platform effect is amplified when generalizing to newly, independently collected cohorts, a platform × evaluation-mode interaction term (LOCOCV vs. independent validation) was fit, adjusting for drug class. This interaction term was not significant (p=0.70), indicating that the magnitude of any platform-associated performance difference does not increase when models are applied to independently collected validation cohorts. Together, these results indicate that rank normalization does not leave a systematic platform bias that would compromise cross-cohort generalizability.

#### Feature comparisons between EXPRESSO models (EXPRESSO-T and EXPRESSO-B)

To evaluate whether the EXPRESSO-Target only model (EXPRESSO-T) captures highly predictive individual genes, we examined the overlap between genes selected by the LASSO-based model and the top-performing gene in univariate analysis. Specifically, we defined top-performing genes as those with mean ROC-AUCs exceeding the 99.9th percentile across all LOCOCV folds. For each fold of the LASSO model, we visualized this overlap using heatmaps, marking top-performing genes selected by LASSO with red stars (Supplementary Figure S6A, S6D, S6G, S6J, S6K, S6L, S6O).

These heatmaps revealed that EXPRESSO-T model frequently selected highly predictive genes in certain contexts—most notably in the trastuzumab and ICB–non-melanoma models. In contrast, the bevacizumab and ICB–melanoma models showed limited overlap. Similarly, for small-molecule targeted therapies and chemotherapies, we again observed limited overlap between LASSO-selected features and top-performing genes. One plausible explanation for this discrepancy is that some of the top univariate genes may be redundant with LASSO, which instead selects correlated alternatives. The EXPRESSO-T model’s limited overlap with the most predictive individual genes motivated the development of the biomarker-augmented model that incorporates differentially expressed genes between responders and non-responders.

From the UpSet plots and Venn diagrams in **Figure S6**, we can observe that in most folds for trastuzumab, and ICB-non-melanoma—where the target gene already demonstrated decent predictive power—additional biomarkers from differentially expressed (DE) genes do not add additional value. For bevacizumab and BRAFi – EXPRESSO-B model was not available as there were no additional biomarkers were available. From **Figure 1C** and **Supplementary Table S3**, we can observe that bevacizumab has the highest clinical heterogeneity index (a weighted index that integrates cancer-type diversity, cohort spread, and treatment mix, details in **Methods**). Whereas, ICB-melanoma has the lowest clinical heterogeneity index, with a single cancer type and monotherapy.

For ICB-melanoma, several non-target genes in the unsupervised setting demonstrated higher predictive value than *CD274* (Figure S6A). The EXPRESSO-T model does not include any of these high-performing features. However, the EXPRESSO-B model—applied to a relatively homogeneous cohort—successfully incorporated additional informative biomarkers in every fold, leading to substantial improvements in both ROC-AUC (from 0.64 to 0.71) and odds ratio (from 1.4 to 4.6). Interestingly, only for ICB-melanoma, each fold identified new biomarkers which were not identified in EXPRESSO-T mode. From the Venn diagram, it is evident that while majority of the features selected in EXPRESSO-B mode are also selected EXPRESSO-T mode, some additional genes are selected in EXPRESSO-B mode improving the prediction accuracy. Several of the genes added by EXPRESSO-B, such as *IRF1*, *UBD*, are known to be biologically relevant for ICB melanoma patients. *IRF1* (identified in five out of nine folds) is a transcription factor that mediates type I and type II interferon responses – related to IFNG signaling and antigen presentation (23,24) – a direct enhancer of ICB responsiveness (25,26). *UBD* is a ubiquitin-like modifier involved in proteasomal degradation and immune activation (Bhat *et al.*, 2022; Hu & Sun, 2016) – whose upregulation may be linked to favorable immune infiltration and ICB responsiveness (29,30).

For BRAF inhibitors, the target gene by itself exhibited poor predictive power. The EXPRESSO-T model successfully captured additional predictive features, boosting the performance. No additional biomarkers are available from differentially expressed (DE) genes from BRAFi cohorts.

For trastuzumab, the target gene *ERBB2* was highly predictive in the unsupervised setting. The EXPRESSO-T model further improved performance by treating *ERBB2* as a regularization-free feature. However, the biomarker-augmented model did not yield further gains. In the feature plot, only two folds identified additional biomarkers, and only one gene is newly identified in biomarker model.

For bevacizumab, data heterogeneity across four cancer types limited the benefit of supervised modeling. The target gene alone performed best in the unsupervised setting, and the supervised EXPRESSO-T model showed marginal improvement in ROC-AUC. No additional biomarkers are available from differentially expressed (DE) genes from bevacizumab cohorts.

For ICB-non-melanoma, which includes five different cancer types and exhibits greater heterogeneity, the EXPRESSO-B model had only four out of 10 folds identifying new genes. Consequently, there was no improvement in predictive performance compared to the EXPRESSO-T model.

For chemotherapies like paclitaxel and FAC-FEC, which lack a clear molecular target and act systemically, identifying cohort-independent biomarkers was particularly challenging. The EXPRESSO-B model failed to generalize, and no additional predictive gains were observed beyond the EXPRESSO-T model.

Overall, these results suggest the importance of both biological relevance and cohort homogeneity when incorporating transcriptomic biomarkers into predictive models. While comparing responders and non-responders can improve performance in certain settings—especially for drugs with limited target signal and low cohort heterogeneity—its utility may be limited in more complex or heterogeneous contexts.

#### EXPRESSO-B biomarker selection and leakage prevention

The EXPRESSO-B biomarker selection procedure is designed with strict data separation to prevent information leakage from the test cohort into the biomarker discovery process. In each outer LOCOCV fold, the test cohort is withheld in its entirety before any analysis begins. Biomarker candidates are identified exclusively from the remaining training cohorts using a nested inner LOCOCV: each inner fold further withholds one training cohort as a validation set, and limma differential expression analysis is performed only on the remaining inner training cohorts. P-values from inner folds are combined using Brown’s method(3) (which accounts for correlation between overlapping training sets), followed by Benjamini–Hochberg FDR(13) correction. Candidate genes must additionally improve mean ROC-AUC by at least 0.02 across inner validation folds (ΔAUC criterion) and reach significance on a Wilcoxon rank-sum test before being included as unpenalized features in the outer fold model.

A consequence of this design is that the biomarker list is specific to each outer fold, because it reflects the differential expression signal and predictive utility observed in the particular set of training cohorts used in that fold. Fold-specific biomarker lists are therefore an expected and intended feature of the method, not an artifact. For ICB-melanoma, this is explicitly illustrated in Supplementary Figure S6A, where each outer fold yields a partially distinct set of biomarkers. The fact that a gene such as CD8A shows significant differential expression in some training cohorts does not guarantee its selection in every fold; selection additionally requires that it consistently improves AUC across inner validation folds of the specific training set used in that outer fold.

Occasional high AUC values in small cohorts (e.g., AUC = 1.0 for GSE123728, n = 13) are consistent with the known statistical behavior of AUC in very small samples, where even a modest score separation can yield a perfect ranking. This is further supported by the positive correlation between training sample size and AUC observed across drugs (Supplementary Figures S3A-G). The complete biomarker selection code has been deposited to Zenodo to allow full independent reproduction of these results.

---

**Figure S1:**
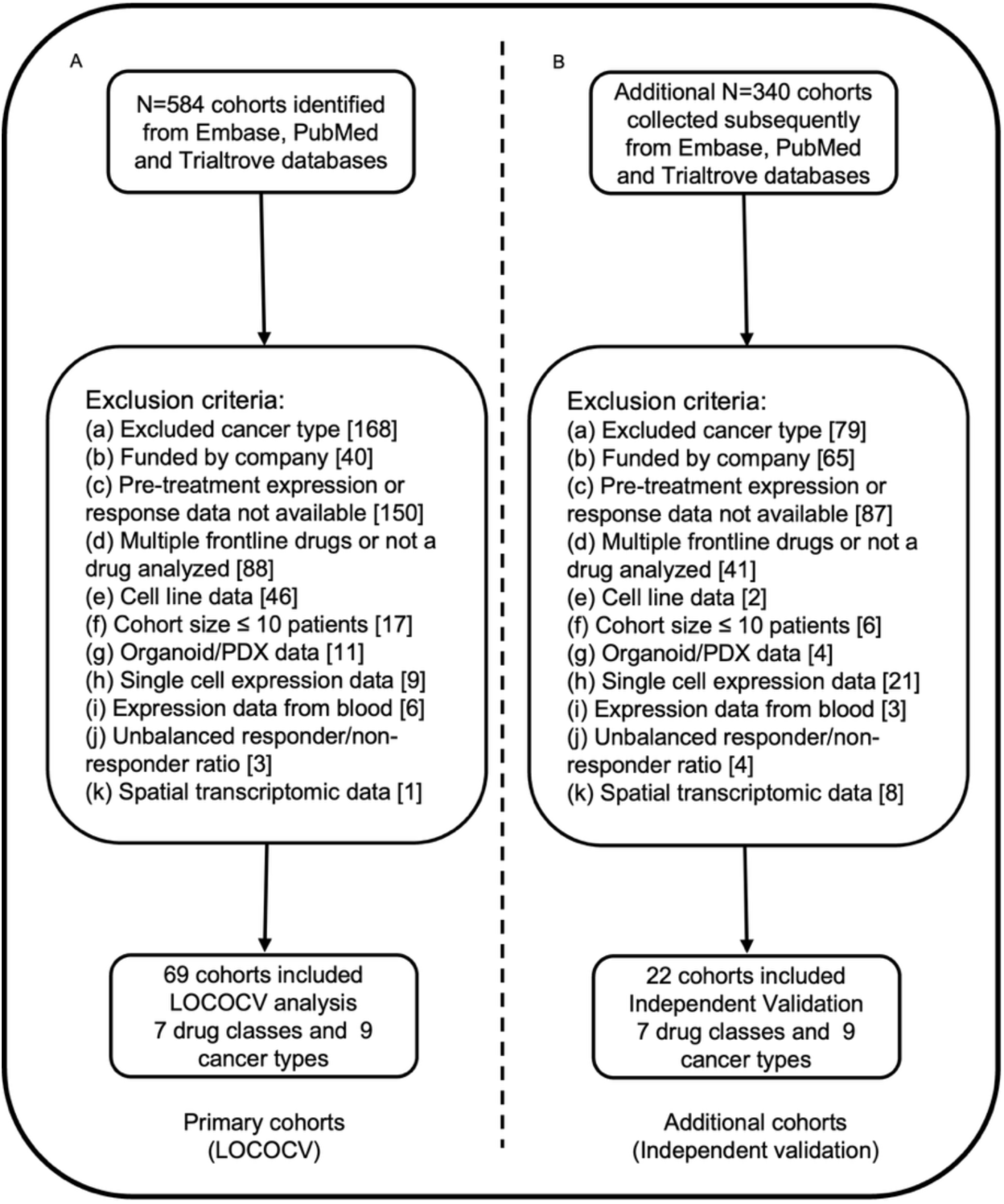
CONSORT-style flowchart summarizing cohort inclusion and exclusion criteria for the primary cohort set used in LOCOCV analysis. (A) and the independent validation cohort set (B). Numbers next to each criterion indicate the count of candidate studies excluded for that reason. When a study was eligible for exclusion under more than one criterion, exclusion was attributed to the first applicable reason encountered when reading the paper, which does not necessarily correspond to the order shown in the diagram, which we tried to follow. The term “CONSORT-style” is used because the items tracked here are research studies, not study participants as in a standard CONSORT diagram.

**Figure S2:**
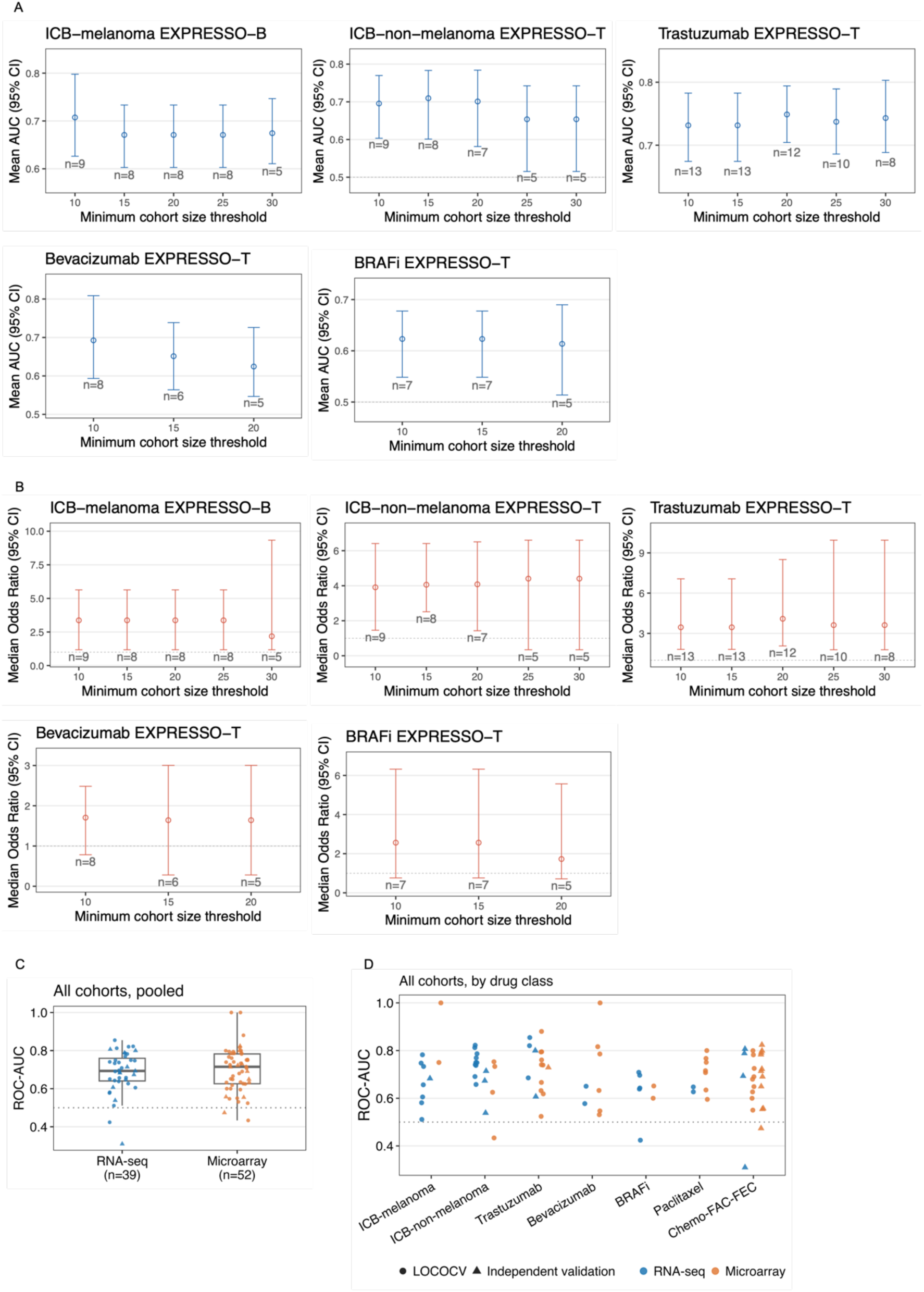
AUC sensitivity analysis. Sensitivity of mean AUC. (A, blue) and median odds ratio (B, red) to minimum cohort size threshold for the ICB-melanoma, ICB-non-melanoma, Trastuzumab, Bevacizumab, and BRAFi analysis. All paclitaxel and chemo-FAC/FEC cohorts exceed 30 patients in size and are therefore not included in this sensitivity analysis. Point estimates (open circles) and 95% confidence intervals (error bars) are shown for thresholds of ≥10 to ≥30 patients in steps of 5; for bevacizumab and BRAFi, thresholds beyond ≥20 are omitted as fewer than 5 cohorts remained, rendering estimates unreliable. The number of cohorts meeting each threshold is indicated (n). Confidence intervals were derived by bootstrap resampling at the cohort level (2,000 iterations, same random seed); where fewer than 5 cohorts remained at a given threshold, a parametric t-distribution was applied instead. Dashed reference lines indicate AUC = 0.5 (chance level) and OR = 1.0 (no effect). Where the number of cohorts retained does not change between consecutive thresholds — because no cohorts have a size falling between the two values — the bootstrap confidence intervals are identical, as the same random seed was applied to the same input values; any difference between adjacent thresholds therefore reflects exclusively the removal of cohorts of size below the higher threshold. Performance estimates remain stable across thresholds, with overlapping confidence intervals throughout; wider intervals at higher thresholds reflect the reduced number of eligible cohorts rather than changes in model performance. (C) Per-cohort ROC-AUC by sequencing platform (RNA-seq, blue; microarray, orange), pooled across all 91 cohorts and both evaluation modes. Box plots show the median and interquartile range; points show individual cohorts, jittered horizontally to reduce overlap, with circles indicating LOCOCV cohorts and triangles indicating independent validation cohorts. The dashed reference line indicates AUC = 0.5 (chance level). Mean AUC did not differ significantly by platform (RNA-seq n=39, mean=0.68; microarray n=52, mean=0.71; Wilcoxon rank-sum test, p=0.50). (D) The same per-cohort AUC values stratified by drug class, with platform indicated by point color and evaluation mode by point shape. After adjusting for drug class in a linear model (AUC ∼ Platform + Drug), the estimated coefficient for platform effect (microarray relative to RNA-seq) remained non-significant (0.039, p=0.19), as it did when weighted by cohort size (p=0.14). A platform × evaluation-mode interaction term testing whether this effect differs between LOCOCV and independent validation was also not significant (p=0.70), indicating no evidence that platform composition affects model generalizability to newly collected cohorts.

**Figure S3:**
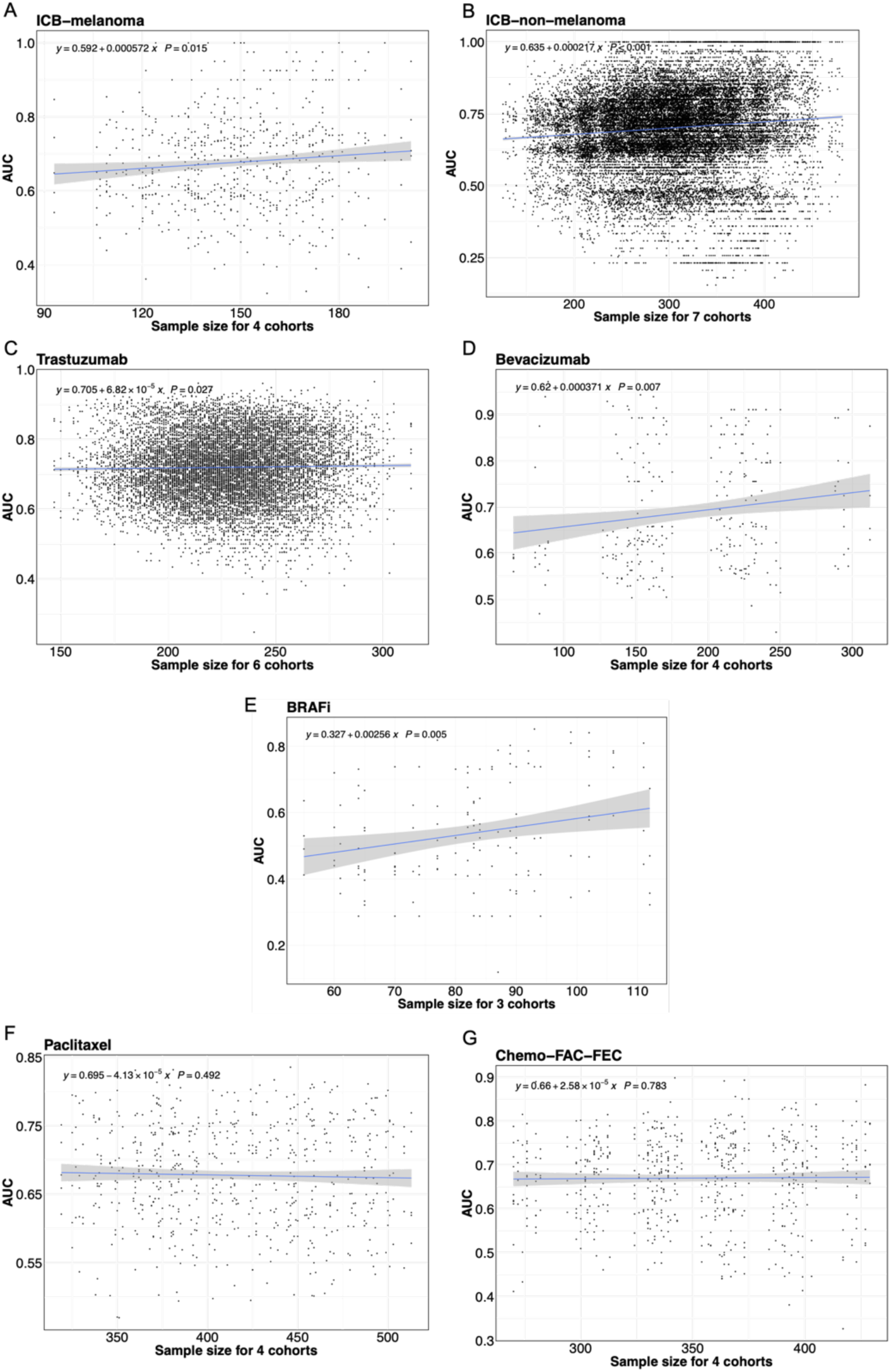
Scatter plots of sample size vs ROC-AUC for fixed numbers of training cohorts. We chose the training cohorts number for each drug with the maximum number of samples.

**Figure S4:**
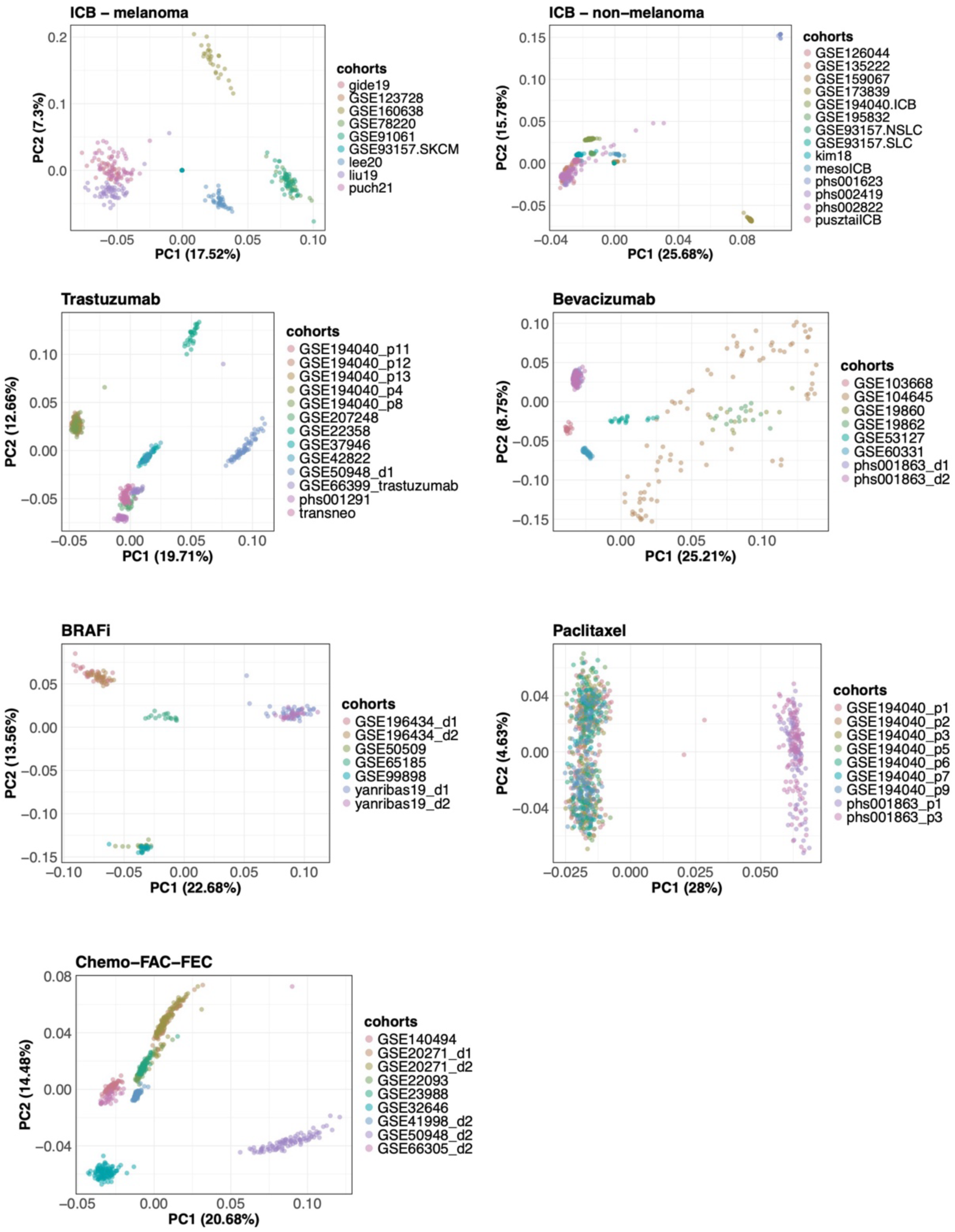
Principal component analysis (PCA) plots of patients with their rank normalized expression data for seven drug-specific cohorts used for EXPRESSO.

**Figure S5:**
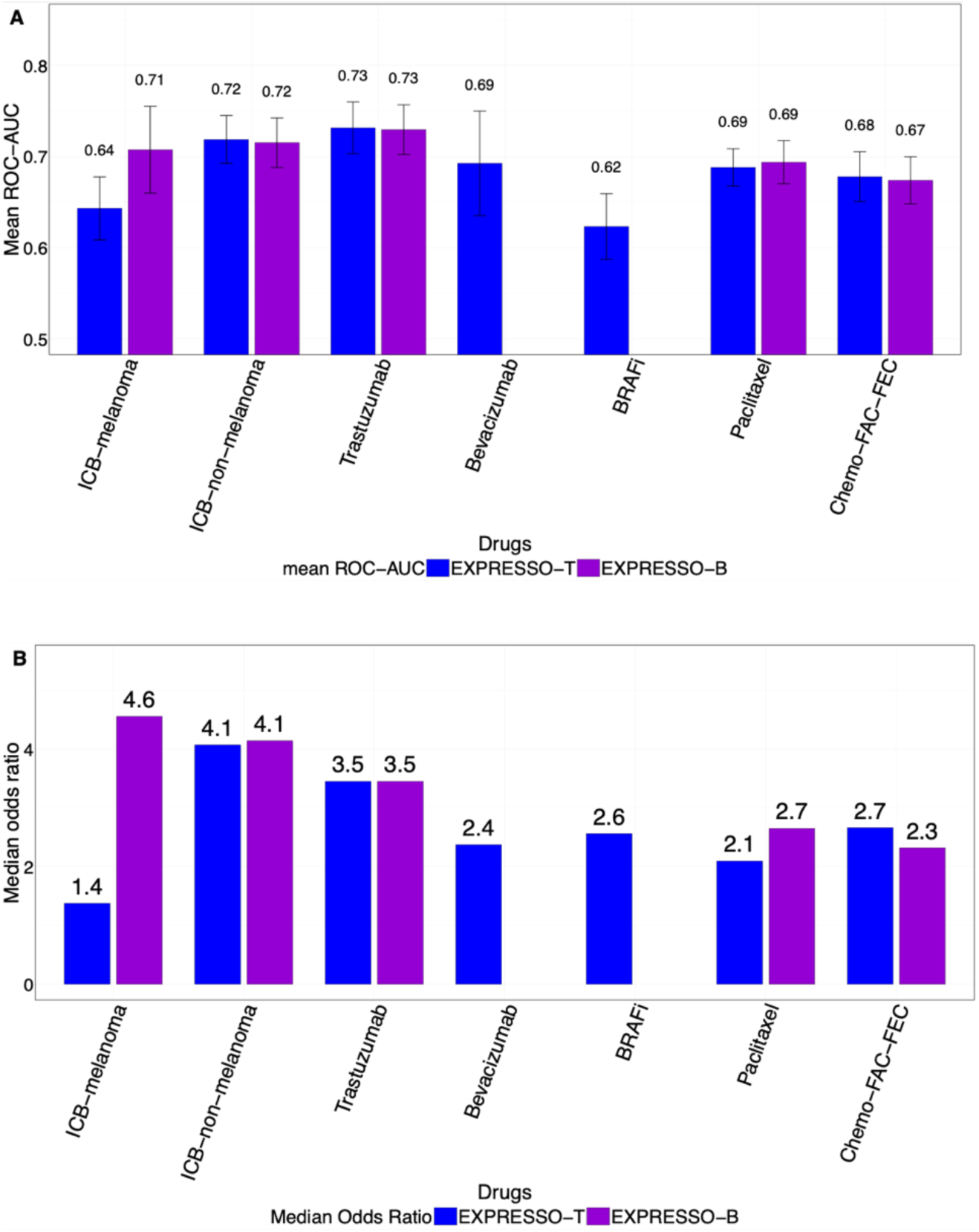
Drug response prediction of EXPRESSO supervised models –. (A) Bar plots for all seven drug-specific supervised EXPRESSO-Target-only models (EXPRESSO-T), compared to EXPRESSO – Target + Biomarker models (EXPRESSO-B) (not available for bevacizumab and BRAFi). (B) Median odds ratios are shown in the bar plot of seven drug-specific EXPRESSO-T compared to EXPRESSO-B models (not available for bevacizumab and BRAFi).

**Figure S6:**
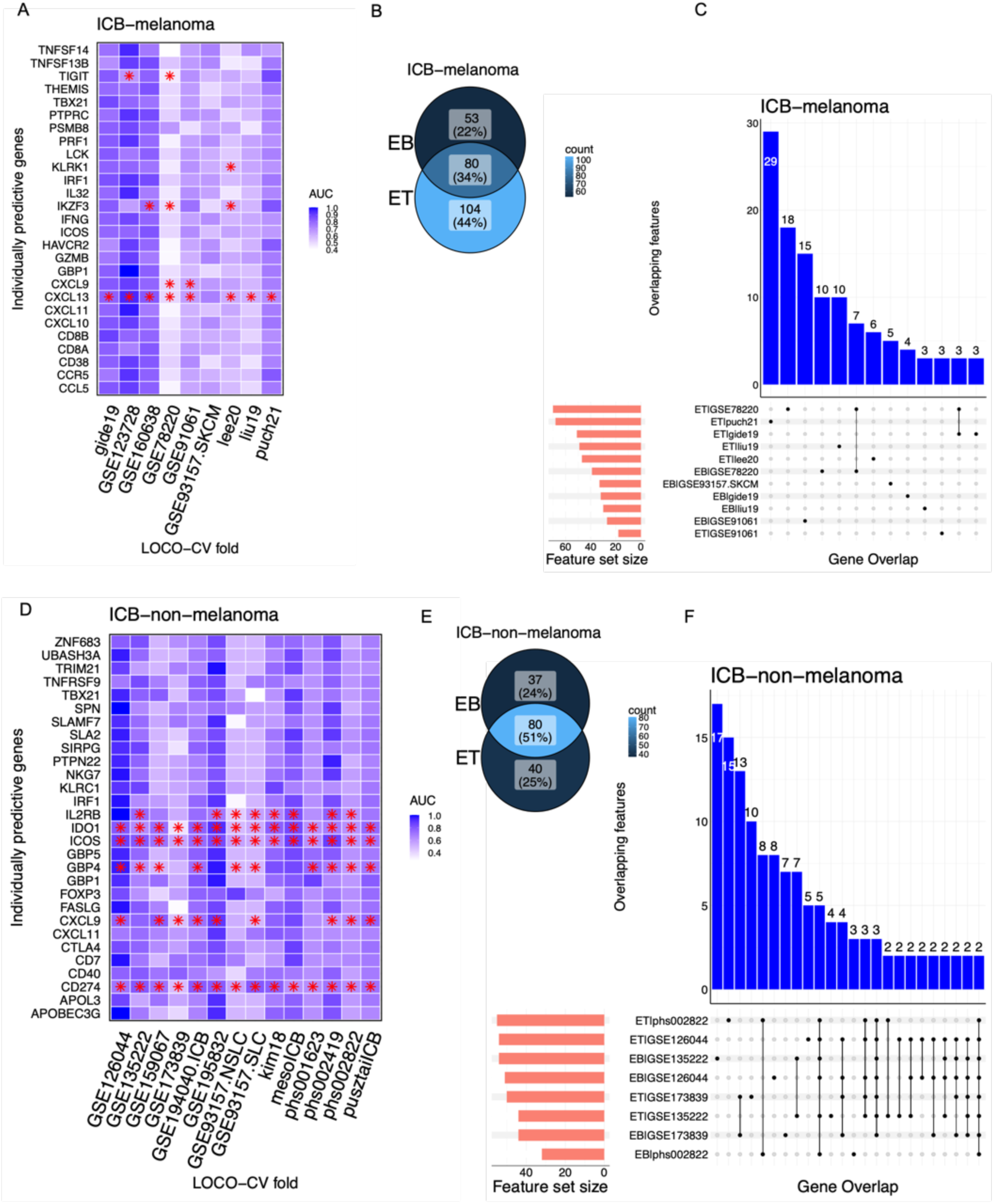
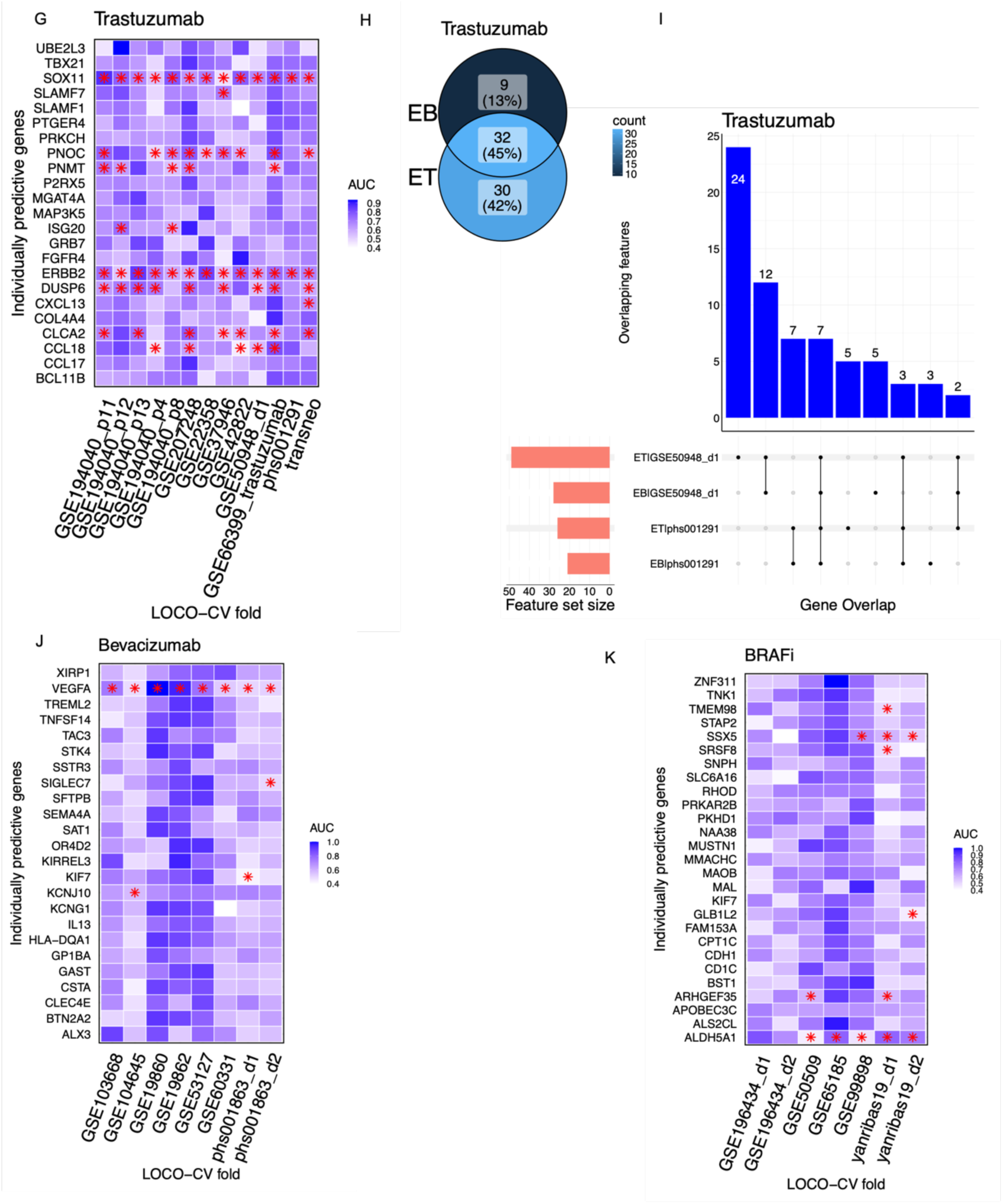
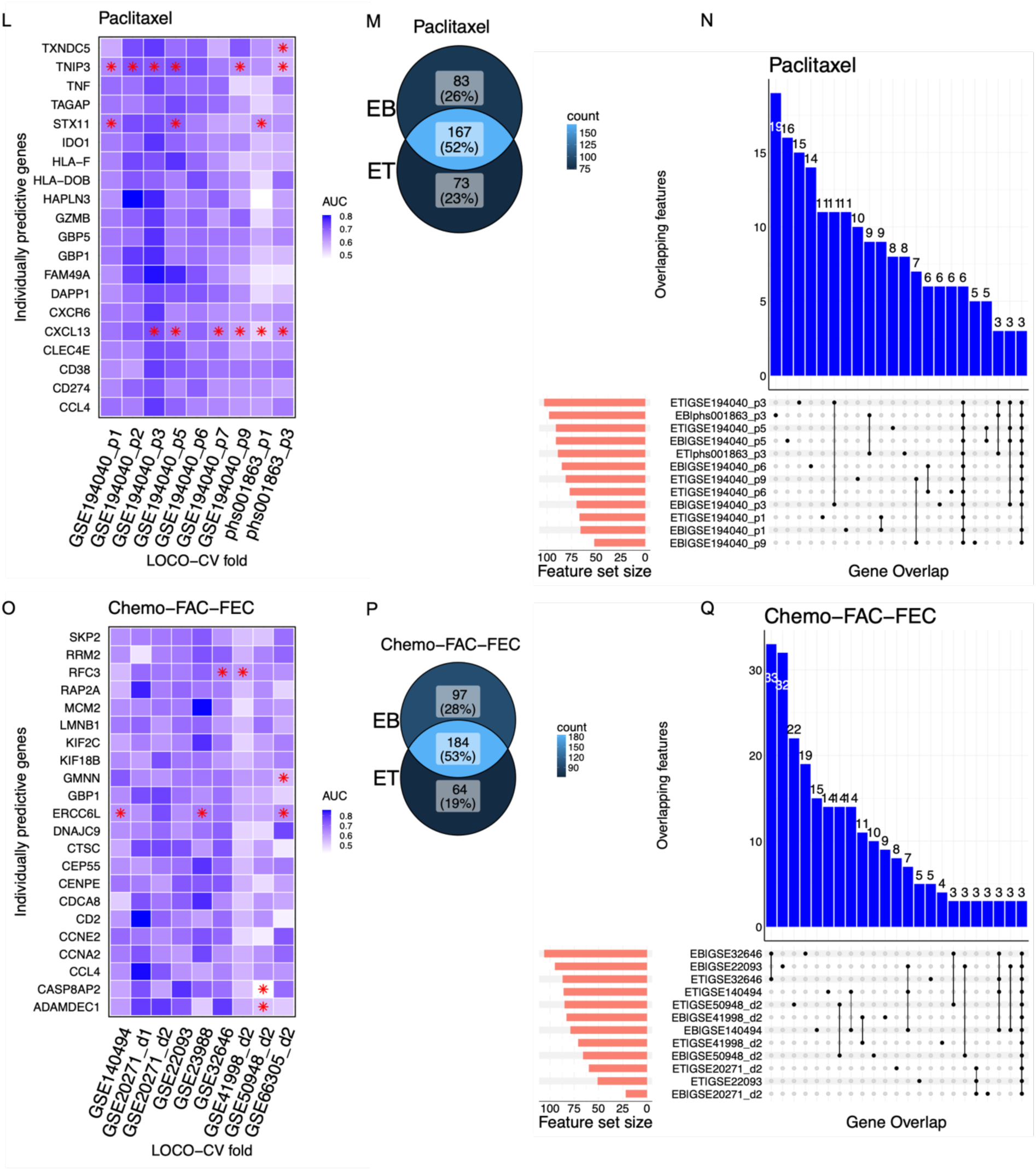
Feature comparisons between models. (A, D, G, J, K, L, O) – Heatmaps show the overlap between genes selected by the LASSO-based EXPRESSO-T model and the top-performing genes identified through unsupervised analysis. We defined top-performing genes as those with mean ROC-AUCs exceeding the 99.9th percentile across all LOCOCV folds for a drug-specific model. For each fold of the LASSO model, we marked top-performing genes selected by LASSO with red stars. (B, E, H, M, P) – Venn-diagrams show the feature overlap between EXPRESSO-T model (ET) and EXPRESSO–B model (EB) (where available, bevacizumab and BRAFi have no EXPRESSO-B model). The lower the counts, the darker the blue shade. (C, F, I, N, Q) – Upset plots show the overlap of gene sets across folds of two supervised models – EXPRESSO-T (ET) and EXPRESSO-B (EB, where available). Horizontal bars at the bottom left show the size of the gene set selected in each fold of the two approaches. Vertical bars at the top show the number of genes shared among the sets indicated by the dot combination directly below. In the dot matrix, each row corresponds to one gene set, labeled on the left as model | cohort ID (e.g., “ET | GSE78220” is the ET model’s gene set from the GSE78220 fold). Within each column, a single filled dot with no connecting line indicates genes uniquely selected in that fold, with no overlap with any other set; dots joined by a line indicate genes shared among the connected sets.

**Figure S7:**
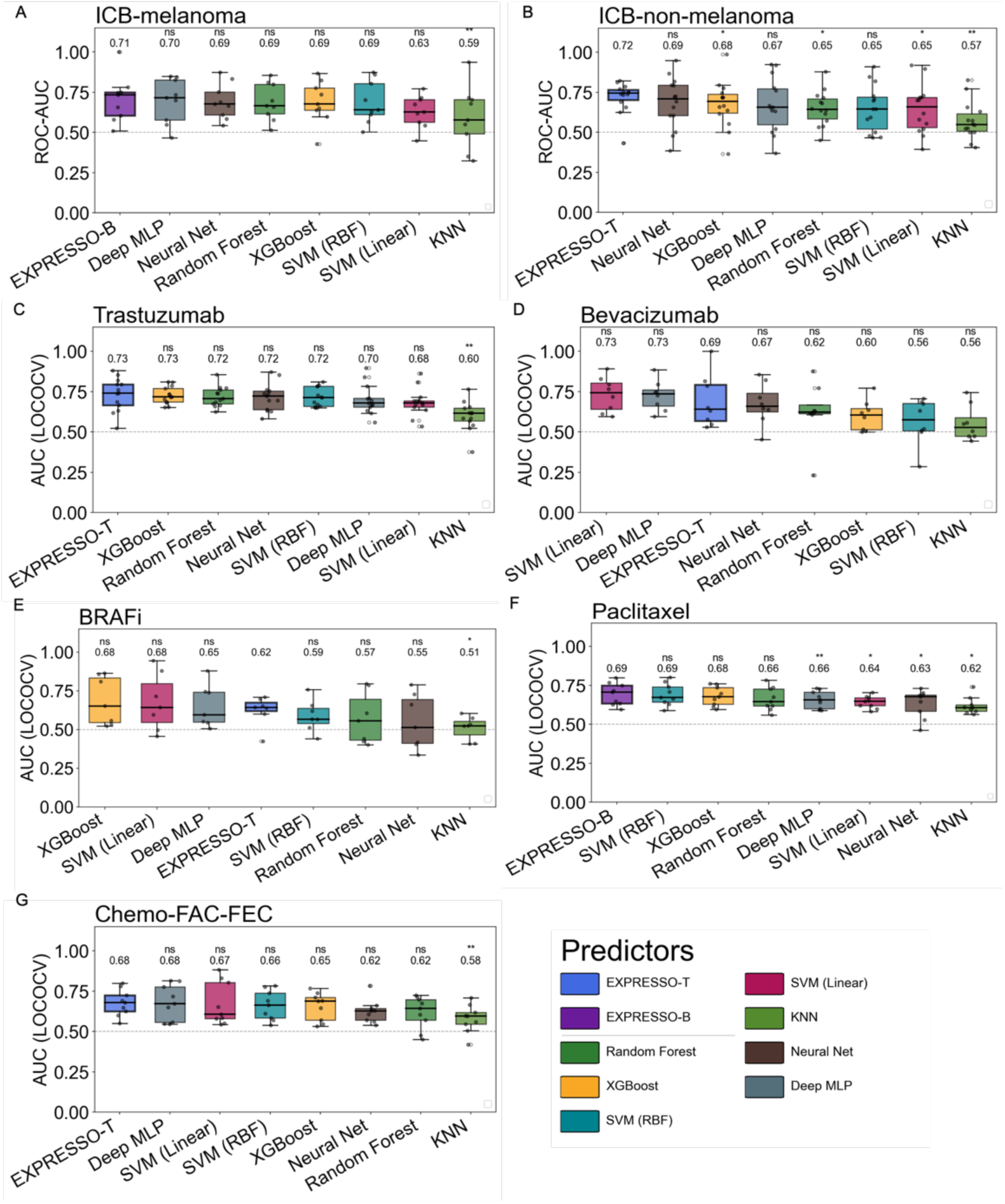
Benchmarking of EXPRESSO against seven machine learning methods across seven drugs under Leave-One-Cohort-Out Cross-Validation (LOCOCV). Boxplots show AUC distribution across independent test cohorts for each method. Mean AUC values are shown above each box. Significance labels indicate two-sided Wilcoxon signed-rank test results versus EXPRESSO (ns: p > 0.05, *: 0.01 ≤ p < 0.05, **: p < 0.01). Dashed line indicates AUC = 0.5 (random classifier).

**Figure S8:**
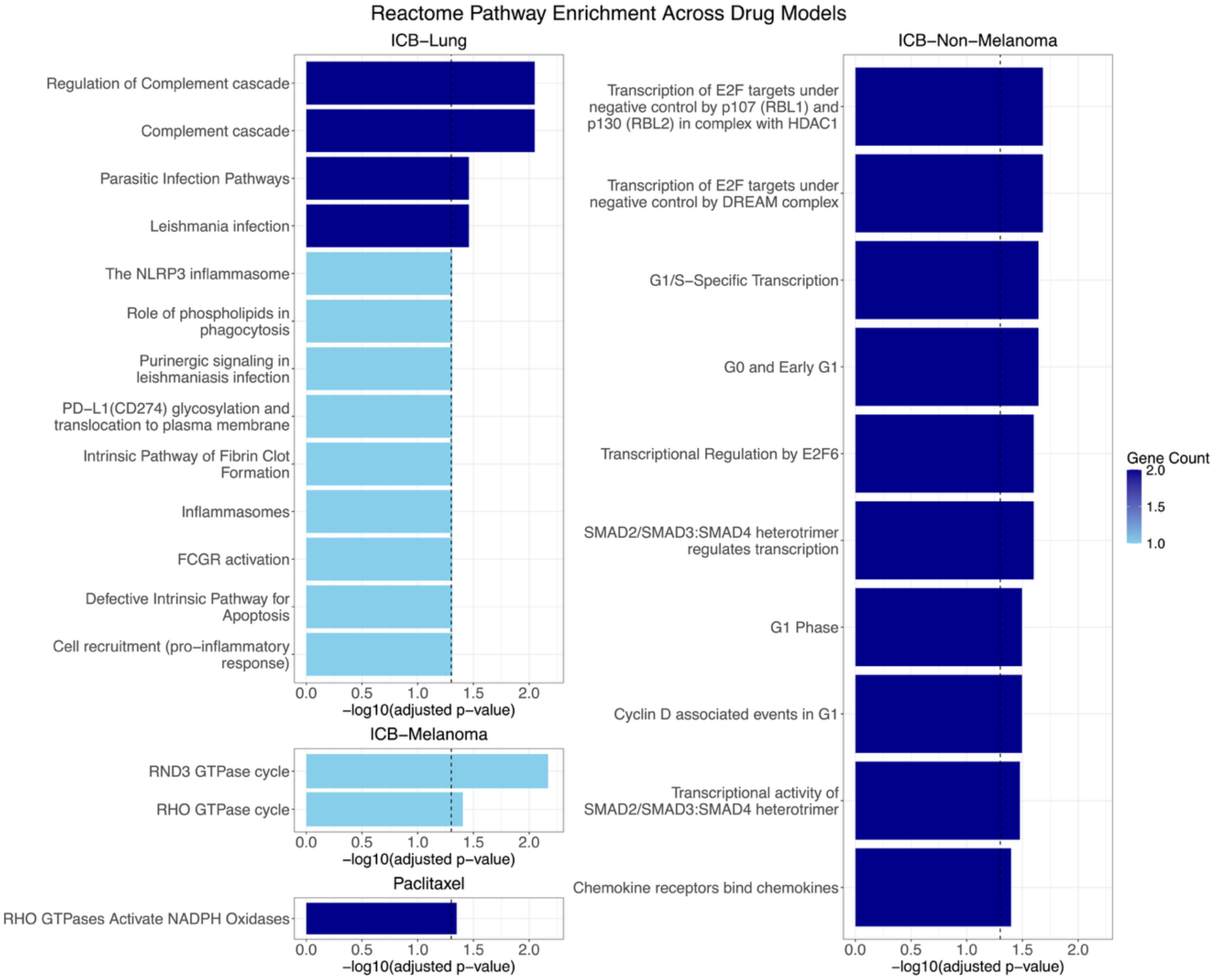
Reactome pathway enrichment analysis of stable model features. Bar plots show significantly enriched Reactome pathways (adjusted p-value <0.05, Benjamini–Hochberg correction) for each drug model with sufficient annotated stable features. The x-axis represents −log10(adjusted p-value) and bar color indicates the number of stable features mapping to each pathway (gene count). The dashed vertical line indicates the significance threshold at p=0.05. Penalty-free target genes were excluded prior to analysis. Models with insufficient mappable annotated features for hypergeometric enrichment testing (trastuzumab, bevacizumab, BRAFi, chemo-FAC-FEC) are not shown.

**Figure S9:**
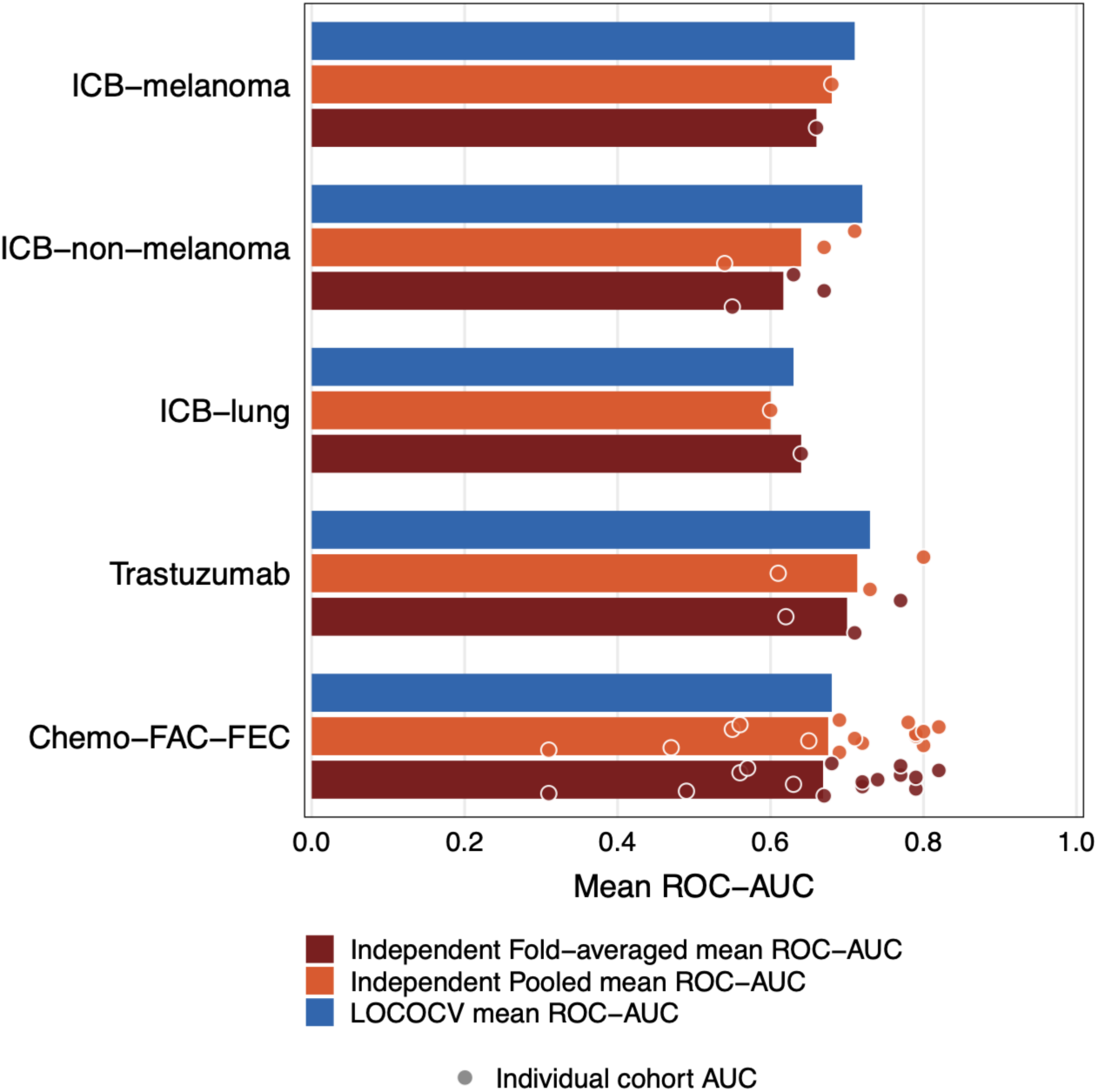
Sensitivity analysis comparing pooled and fold-averaged independent validation approaches. Horizontal bars show mean ROC-AUC for each drug category under three evaluation strategies: LOCOCV (blue), independent validation using a pooled model trained on all original cohorts (orange), and independent validation using a fold-averaged approach, in which each held-out cohort was tested against every LOCOCV fold model and ROC-AUCs were averaged across folds (maroon). Individual cohort ROC-AUCs are overlaid as solid circles with white borders on their respective independent validation bars. Independent cohorts were available for five drug categories: ICB-melanoma (1 cohort), ICB-non-melanoma (3 cohorts), ICB-lung (1 cohort), trastuzumab (3 cohorts), and chemo-FAC-FEC (15 cohorts). The pooled and fold-averaged approaches yield comparable performance across all drug categories, indicating that EXPRESSO generalizes stably regardless of the evaluation strategy.

**Figure S10:**
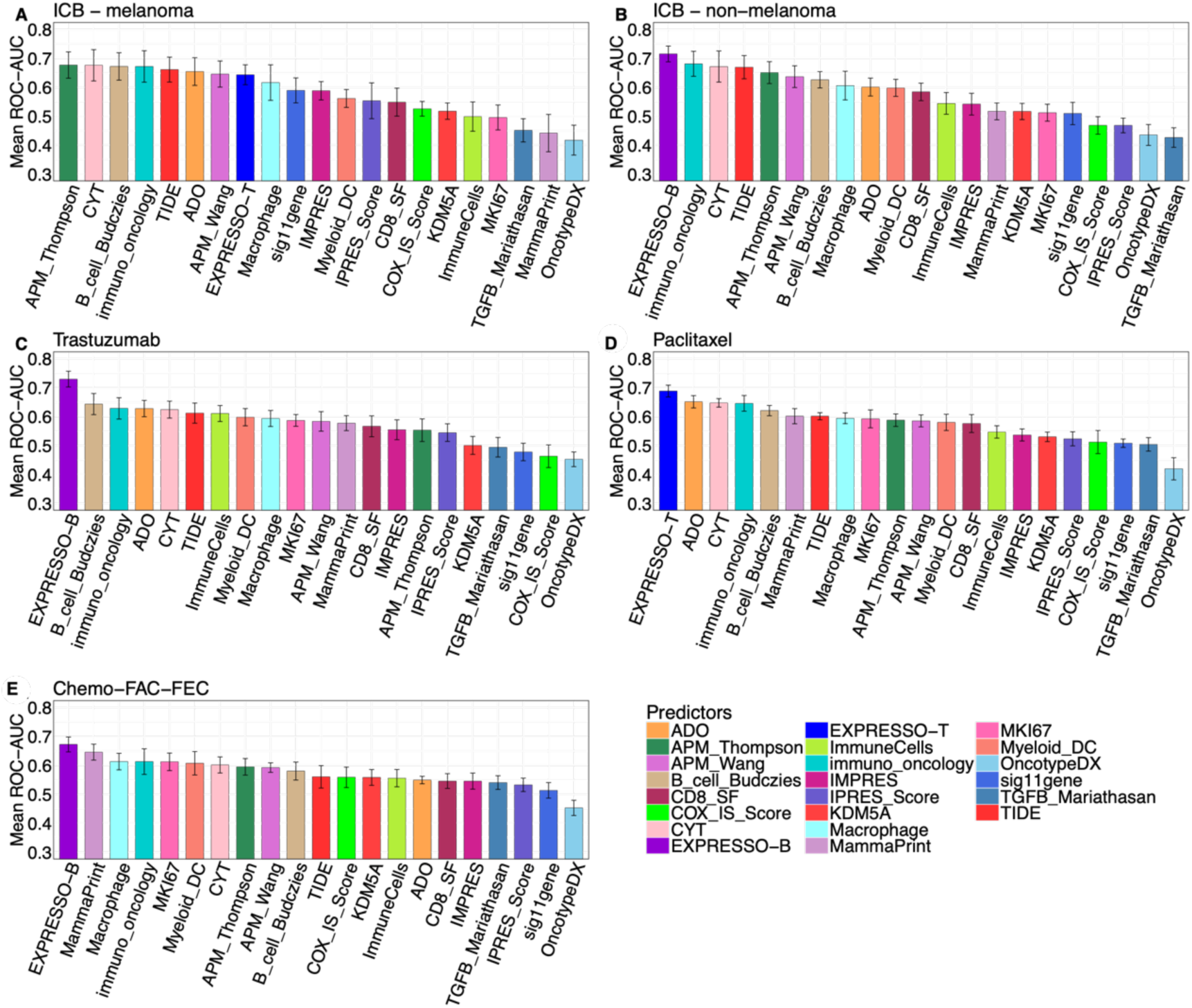
Comparative performance of worst-performing EXPRESSO variant for each drug class against existing gene signatures and methods. Bar plots show the mean ROC-AUC values for each predictor across drug-specific models, ordered from highest to lowest mean performance. While many of the published signatures were originally developed for specific therapeutic or disease contexts, we applied the same set of 20 signatures uniformly across all drugs to enable consistent benchmarking. The worst-performing EXPRESSO variant (EXPRESSO-T or EXPRESSO-B) is shown for each drug (except bevacizumab and BRAFi, where EXPRESSO-B model is not available), while the best performing EXPRESSO variant for each drug is shown in Figure 5.

**Figure S11:**
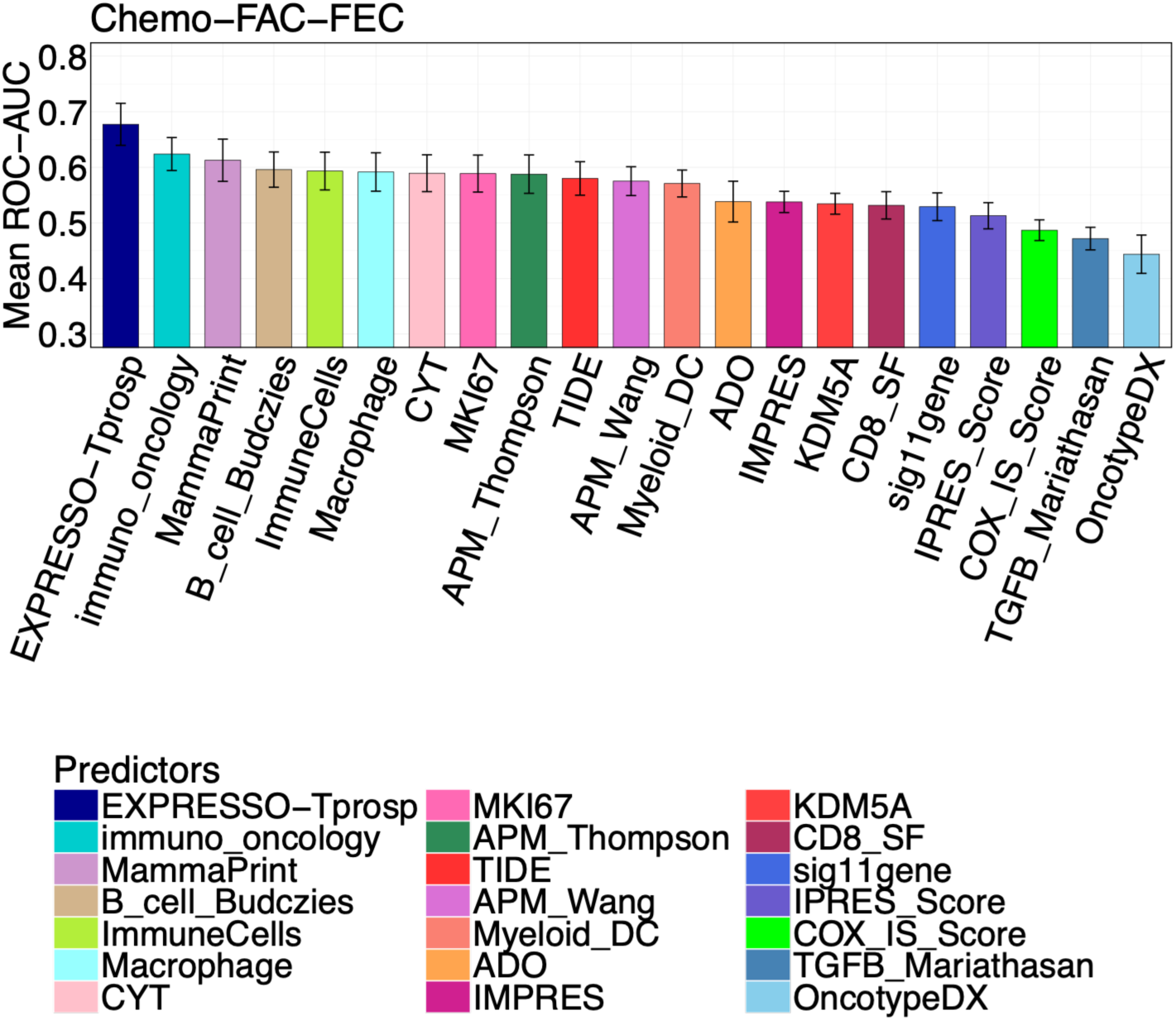
Benchmarking EXPRESSO against published transcriptomic signatures on independent Chemo-FAC-FEC cohorts. Mean ROC-AUC of EXPRESSO-Tprosp (pooled model) and 20 published transcriptomic signatures evaluated on 15 independently held-out Chemo-FAC-FEC cohorts that were not used during model development. EXPRESSO-Tprosp is highlighted as the top-performing predictor. This comparison was performed for Chemo-FAC-FEC only, as it was the sole drug category with enough number of independent cohorts (n=15) to support a reliable multi-method benchmark.

## REFERENCES

1. PCAWG Transcriptome Core Group, Calabrese C, Davidson NR, Demircioğlu D, Fonseca NA, He Y, et al. Genomic basis for RNA alterations in cancer. Nature. 2020;578:129–36.

2. He X, Xu C. Immune checkpoint signaling and cancer immunotherapy. Cell Res. 2020;30:660–9.

3. Justiz-Vaillant A, Pandit BR, Unakal C, Vuma S, Akpaka PE. A comprehensive review about the use of monoclonal antibodies in cancer therapy. Antibodies Basel Switz. 2025;14:35.

4. Liu G-H, Chen T, Zhang X, Ma X-L, Shi H-S. Small molecule inhibitors targeting the cancers. MedComm. 2022;3:e181.

5. Nesic K, Parker P, Swisher EM, Krais JJ. DNA repair and the contribution to chemotherapy resistance. Genome Med. 2025;17:62.

6. Tsimberidou AM, Fountzilas E, Bleris L, Kurzrock R. Transcriptomics and solid tumors: the next frontier in precision cancer medicine. Semin Cancer Biol. 2022;84:50–9.

7. Hoadley KA, Yau C, Wolf DM, Cherniack AD, Tamborero D, Ng S, et al. Multiplatform analysis of 12 cancer types reveals molecular classification within and across tissues of origin. Cell. 2014;158:929–44.

8. Passaro A, Al Bakir M, Hamilton EG, Diehn M, André F, Roy-Chowdhuri S, et al. Cancer biomarkers: emerging trends and clinical implications for personalized treatment. Cell. 2024;187:1617–35.

9. Chakravarty D, Gao J, Phillips S, Kundra R, Zhang H, Wang J, et al. OncoKB: A Precision Oncology Knowledge Base. JCO Precis Oncol. 2017;1–16.

10. Li D, Chan DW. Proteomic cancer biomarkers from discovery to approval: it’s worth the effort. Expert Rev Proteomics. 2014;11:135–6.

11. Garon EB, Rizvi NA, Hui R, Leighl N, Balmanoukian AS, Eder JP, et al. Pembrolizumab for the treatment of non-small-cell lung cancer. N Engl J Med. 2015;372:2018–28.

12. Sparano JA, Gray RJ, Makower DF, Pritchard KI, Albain KS, Hayes DF, et al. Adjuvant chemotherapy guided by a 21-gene expression assay in breast cancer. N Engl J Med. 2018;379:111–21.

13. Ayers M, Lunceford J, Nebozhyn M, Murphy E, Loboda A, Kaufman DR, et al. IFN-γ-related mRNA profile predicts clinical response to PD-1 blockade. J Clin Invest. 2017;127:2930–40.

14. Dinstag G, Shulman ED, Elis E, Ben-Zvi DS, Tirosh O, Maimon E, et al. Clinically oriented prediction of patient response to targeted and immunotherapies from the tumor transcriptome. Med N Y N. 2023;4:15–30.e8.

15. Geeleher P, Cox NJ, Huang RS. Clinical drug response can be predicted using baseline gene expression levels and in vitro drug sensitivity in cell lines. Genome Biol. 2014;15:R47.

16. Lee J-K, Liu Z, Sa JK, Shin S, Wang J, Bordyuh M, et al. Pharmacogenomic landscape of patient-derived tumor cells informs precision oncology therapy. Nat Genet. 2018;50:1399–411.

17. Warren A, Chen Y, Jones A, Shibue T, Hahn WC, Boehm JS, et al. Global computational alignment of tumor and cell line transcriptional profiles. Nat Commun. 2021;12:22.

18. Ma J, Fong SH, Luo Y, Bakkenist CJ, Shen JP, Mourragui S, et al. Few-shot learning creates predictive models of drug response that translate from high-throughput screens to individual patients. Nat Cancer. 2021;2:233–44.

19. He D, Liu Q, Wu Y, Xie L. A context-aware deconfounding autoencoder for robust prediction of personalized clinical drug response from cell-line compound screening. Nat Mach Intell. 2022;4:879–92.

20. Kuenzi BM, Park J, Fong SH, Sanchez KS, Lee J, Kreisberg JF, et al. Predicting drug response and synergy using a deep learning model of human cancer cells. Cancer Cell. 2020;38:672–684.e6.

21. Partin A, Brettin TS, Zhu Y, Narykov O, Clyde A, Overbeek J, et al. Deep learning methods for drug response prediction in cancer: predominant and emerging trends. Front Med. 2023;10:1086097.

22. Shen W, Nguyen TH, Li MM, Huang Y, Moon I, Nair N, et al. Generalizable AI predicts immunotherapy outcomes across cancers and treatments [Internet]. Pharmacology and Therapeutics; 2025 [cited 2025 Sept 24]. Available from: http://medrxiv.org/lookup/doi/10.1101/2025.05.01.25326820

23. Ling A, Huang RS. Computationally predicting clinical drug combination efficacy with cancer cell line screens and independent drug action. Nat Commun. 2020;11:5848.

24. Beaubier N, Bontrager M, Huether R, Igartua C, Lau D, Tell R, et al. Integrated genomic profiling expands clinical options for patients with cancer. Nat Biotechnol. 2019;37:1351–60.

25. Ianevski A, Nader K, Driva K, Senkowski W, Bulanova D, Moyano-Galceran L, et al. Single-cell transcriptomes identify patient-tailored therapies for selective co-inhibition of cancer clones. Nat Commun. 2024;15:8579.

26. Pleasance E, Bohm A, Williamson LM, Nelson JMT, Shen Y, Bonakdar M, et al. Whole-genome and transcriptome analysis enhances precision cancer treatment options. Ann Oncol. 2022;33:939–49.

27. Mundi PS, Dela Cruz FS, Grunn A, Diolaiti D, Mauguen A, Rainey AR, et al. A transcriptome-based precision oncology platform for patient-therapy alignment in a diverse set of treatment-resistant malignancies. Cancer Discov. 2023;13:1386–407.

28. Irmisch A, Bonilla X, Chevrier S, Lehmann K-V, Singer F, Toussaint NC, et al. The tumor profiler study: integrated, multi-omic, functional tumor profiling for clinical decision support. Cancer Cell. 2021;39:288–93.

29. Borisov N, Buzdin A. Transcriptomic harmonization as the way for suppressing cross-platform bias and batch effect. Biomedicines. 2022;10:2318.

30. Feng J, Ren J, Yang Q, Liao L, Cui L, Gong Y, et al. Metabolic gene signature for predicting breast cancer recurrence using transcriptome analysis. Future Oncol. 2021;17:71–80.

31. Wu Y, Liu Y, He A, Guan B, He S, Zhang C, et al. Identification of the six-RNA-binding protein signature for prognosis prediction in bladder cancer. Front Genet. 2020;11:992.

32. Parker HS, Corrada Bravo H, Leek JT. Removing batch effects for prediction problems with frozen surrogate variable analysis. PeerJ. 2014;2:e561.

33. Clough E, Barrett T, Wilhite SE, Ledoux P, Evangelista C, Kim IF, et al. NCBI GEO: archive for gene expression and epigenomics data sets: 23-year update. Nucleic Acids Res. 2024;52:D138–44.

34. Langfelder P, Horvath S. WGCNA: an R package for weighted correlation network analysis. BMC Bioinformatics. 2008;9:559.

35. Tryka KA, Hao L, Sturcke A, Jin Y, Wang ZY, Ziyabari L, et al. NCBI’s database of genotypes and phenotypes: dbGaP. Nucleic Acids Res. 2014;42:D975–9.

36. Patro R, Duggal G, Love MI, Irizarry RA, Kingsford C. Salmon provides fast and bias-aware quantification of transcript expression. Nat Methods. 2017;14:417–9.

37. Soneson C, Love MI, Robinson MD. Differential analyses for RNA-seq: transcript-level estimates improve gene-level inferences. F1000Research. 2015;4:1521.

38. Friedman J, Hastie T, Tibshirani R. Regularization paths for generalized linear models via coordinate descent. J Stat Softw. 2010;33:1–22.

39. Kassambara A, Kosinski M, Biecek P. survminer: Drawing Survival Curves using “ggplot2”. [Internet]. Available from: https://rpkgs.datanovia.com/survminer/index.html

40. De Sousa Linhares A, Battin C, Jutz S, Leitner J, Hafner C, Tobias J, et al. Therapeutic PD-L1 antibodies are more effective than PD-1 antibodies in blocking PD-1/PD-L1 signaling. Sci Rep. 2019;9:11472.

41. Castellani G, Buccarelli M, Arasi MB, Rossi S, Pisanu ME, Bellenghi M, et al. BRAF mutations in melanoma: biological aspects, therapeutic implications, and circulating biomarkers. Cancers. 2023;15:4026.

42. Jordan MA, Wilson L. Microtubules as a target for anticancer drugs. Nat Rev Cancer. 2004;4:253–65.

43. Marinello J, Delcuratolo M, Capranico G. Anthracyclines as topoisomerase II poisons: from early studies to new perspectives. Int J Mol Sci. 2018;19:3480.

44. Kuremsky JG, Tepper JE, McLeod HL. Biomarkers for response to neoadjuvant chemoradiation for rectal cancer. Int J Radiat Oncol Biol Phys. 2009;74:673–88.

45. Dubacheva GV, Curk T, Richter RP. Determinants of superselectivity─practical concepts for application in biology and medicine. Acc Chem Res. 2023;56:729–39.

46. Purbey PK, Seo J, Paul MK, Iwamoto KS, Daly AE, Feng A-C, et al. Opposing tumor-cell-intrinsic and –extrinsic roles of the IRF1 transcription factor in antitumor immunity. Cell Rep. 2024;43:114289.

47. Hu X, Wang J, Chu M, Liu Y, Wang Z-W, Zhu X. Emerging role of ubiquitination in the regulation of PD-1/PD-L1 in cancer immunotherapy. Mol Ther J Am Soc Gene Ther. 2021;29:908–19.

48. Newman AM, Liu CL, Green MR, Gentles AJ, Feng W, Xu Y, et al. Robust enumeration of cell subsets from tissue expression profiles. Nat Methods. 2015;12:453–7.

49. Sidders B, Zhang P, Goodwin K, O’Connor G, Russell DL, Borodovsky A, et al. Adenosine signaling is prognostic for cancer outcome and has predictive utility for immunotherapeutic response. Clin Cancer Res. 2020;26:2176–87.

50. Helmink BA, Reddy SM, Gao J, Zhang S, Basar R, Thakur R, et al. B cells and tertiary lymphoid structures promote immunotherapy response. Nature. 2020;577:549–55.

51. Mariathasan S, Turley SJ, Nickles D, Castiglioni A, Yuen K, Wang Y, et al. TGFβ attenuates tumour response to PD-L1 blockade by contributing to exclusion of T cells. Nature. 2018;554:544–8.

52. Thompson JC, Davis C, Deshpande C, Hwang W-T, Jeffries S, Huang A, et al. Gene signature of antigen processing and presentation machinery predicts response to checkpoint blockade in non-small cell lung cancer (NSCLC) and melanoma. J Immunother Cancer. 2020;8:e000974.

53. Wang S, He Z, Wang X, Li H, Liu X-S. Antigen presentation and tumor immunogenicity in cancer immunotherapy response prediction. eLife. 2019;8:e49020.

54. Budczies J, Kirchner M, Kluck K, Kazdal D, Glade J, Allgäuer M, et al. A gene expression signature associated with B cells predicts benefit from immune checkpoint blockade in lung adenocarcinoma. Oncoimmunology. 2021;10:1860586.

55. Sade-Feldman M, Yizhak K, Bjorgaard SL, Ray JP, de Boer CG, Jenkins RW, et al. Defining T cell states associated with response to checkpoint immunotherapy in melanoma. Cell. 2018;175:998–1013.e20.

56. Rooney MS, Shukla SA, Wu CJ, Getz G, Hacohen N. Molecular and genetic properties of tumors associated with local immune cytolytic activity. Cell. 2015;160:48–61.

57. Xiong D, Wang Y, You M. A gene expression signature of TREM2hi macrophages and γδ T cells predicts immunotherapy response. Nat Commun. 2020;11:5084.

58. Auslander N, Zhang G, Lee JS, Frederick DT, Miao B, Moll T, et al. Robust prediction of response to immune checkpoint blockade therapy in metastatic melanoma. Nat Med. 2018;24:1545–9.

59. Hugo W, Shi H, Sun L, Piva M, Song C, Kong X, et al. Non-genomic and immune evolution of melanoma acquiring MAPKi resistance. Cell. 2015;162:1271–85.

60. Bonavita E, Bromley CP, Jonsson G, Pelly VS, Sahoo S, Walwyn-Brown K, et al. Antagonistic inflammatory phenotypes dictate tumor fate and response to immune checkpoint blockade. Immunity. 2020;53:1215–1229.e8.

61. Jiang P, Gu S, Pan D, Fu J, Sahu A, Hu X, et al. Signatures of T cell dysfunction and exclusion predict cancer immunotherapy response. Nat Med. 2018;24:1550–8.

62. Wang L, Gao Y, Zhang G, Li D, Wang Z, Zhang J, et al. Enhancing KDM5A and TLR activity improves the response to immune checkpoint blockade. Sci Transl Med. 2020;12:eaax2282.

63. Davey MG, Hynes SO, Kerin MJ, Miller N, Lowery AJ. Ki-67 as a prognostic biomarker in invasive breast cancer. Cancers. 2021;13:4455.

64. Slodkowska EA, Ross JS. MammaPrint 70-gene signature: another milestone in personalized medical care for breast cancer patients. Expert Rev Mol Diagn. 2009;9:417–22.

65. Syed YY. Oncotype DX breast recurrence score®: a review of its use in early-stage breast cancer. Mol Diagn Ther. 2020;24:621–32.

66. Iwase T, Blenman KRM, Li X, Reisenbichler E, Seitz R, Hout D, et al. A novel immunomodulatory 27-gene signature to predict response to neoadjuvant immunochemotherapy for primary triple-negative breast cancer. Cancers. 2021;13:4839.

67. Shen J, Yan D, Bai L, Geng R, Zhao X, Li H, et al. An 11-gene signature based on treatment responsiveness predicts radiation therapy survival benefit among breast cancer patients. Front Oncol. 2021;11:816053.

68. Wu M, Huang Q, Xie Y, Wu X, Ma H, Zhang Y, et al. Improvement of the anticancer efficacy of PD-1/PD-L1 blockade via combination therapy and PD-L1 regulation. J Hematol OncolJ Hematol Oncol. 2022;15:24.

69. Lin A, Giuliano CJ, Palladino A, John KM, Abramowicz C, Yuan ML, et al. Off-target toxicity is a common mechanism of action of cancer drugs undergoing clinical trials. Sci Transl Med. 2019;11:eaaw8412.

70. Kim Y, Nagy M, Pollard B, Rajagopal PS. Optimizing transcriptome-based synthetic lethality predictions to improve precision oncology in early-stage breast cancer: BC-SELECT [Internet]. Bioinformatics; 2024 [cited 2024 Dec 4]. Available from: http://biorxiv.org/lookup/doi/10.1101/2024.08.15.608073

71. Banna GL, Cantale O, Bersanelli M, Del Re M, Friedlaender A, Cortellini A, et al. Are anti-PD1 and anti-PD-L1 alike? The non-small-cell lung cancer paradigm. Oncol Rev. 2020;14:490.

72. Hoang D-T, Dinstag G, Shulman ED, Hermida LC, Ben-Zvi DS, Elis E, et al. A deep-learning framework to predict cancer treatment response from histopathology images through imputed transcriptomics. Nat Cancer. 2024;5:1305–17.

73. Hoang D-T, Shulman ED, Dhruba SR, Nair NU, Barman RK, Lalchungnunga H, et al. Path2Omics: Enhanced transcriptomic and methylation prediction accuracy from tumor histopathology. BioRxiv Prepr Serv Biol. 2025;2025.02.26.640189.

74. Shulman ED, Campagnolo EM, Lodha R, Stemmer A, Cantore T, Ru B, et al. AI-Driven Spatial Transcriptomics Unlocks Large-Scale Breast Cancer Biomarker Discovery from Histopathology [Internet]. Cancer Biology; 2024 [cited 2025 Sept 26]. Available from: http://biorxiv.org/lookup/doi/10.1101/2024.10.16.618609

75. Singhal A, Zhao X, Wall P, So E, Calderini G, Partin A, et al. The Hallmarks of Predictive Oncology. Cancer Discov. 2025;15:271–85.

## REFERENCES

1. Friedman J, Hastie T, Tibshirani R. Regularization paths for generalized linear models via coordinate descent. J Stat Softw. 2010;33:1–22.

2. Ritchie ME, Phipson B, Wu D, Hu Y, Law CW, Shi W, et al. limma powers differential expression analyses for RNA-sequencing and microarray studies. Nucleic Acids Res. 2015;43:e47.

3. Poole W, Gibbs DL, Shmulevich I, Bernard B, Knijnenburg TA. Combining dependent P-values with an empirical adaptation of Brown’s method. Bioinformatics. Oxford, England; 2016;32:i430–6.

4. Jiang P, Gu S, Pan D, Fu J, Sahu A, Hu X, et al. Signatures of T cell dysfunction and exclusion predict cancer immunotherapy response. Nat Med. 2018;24:1550–8.

5. Auslander N, Zhang G, Lee JS, Frederick DT, Miao B, Moll T, et al. Robust prediction of response to immune checkpoint blockade therapy in metastatic melanoma. Nat Med. 2018;24:1545–9.

6. Hugo W, Zaretsky JM, Sun L, Song C, Moreno BH, Hu-Lieskovan S, et al. Genomic and transcriptomic features of response to anti-PD-1 therapy in metastatic melanoma. Cell. 2016;165:35–44.

7. Bonavita E, Bromley CP, Jonsson G, Pelly VS, Sahoo S, Walwyn-Brown K, et al. Antagonistic inflammatory phenotypes dictate tumor fate and response to immune checkpoint blockade. Immunity. 2020;53:1215–1229.e8.

8. Abraham A, Pedregosa F, Eickenberg M, Gervais P, Mueller A, Kossaifi J, et al. Machine learning for neuroimaging with scikit-learn. Front Neuroinformatics. 2014;8:14.

9. Chen T, Guestrin C. XGBoost: A Scalable Tree Boosting System. Proc 22nd ACM SIGKDD Int Conf Knowl Discov Data Min [Internet]. San Francisco California USA: ACM; 2016 [cited 2026 May 18]. page 785–94. Available from: https://dl.acm.org/doi/10.1145/2939672.2939785

10. Paszke A, Gross S, Massa F, Lerer A, Bradbury J, Chanan G, et al. PyTorch: An Imperative Style, High-Performance Deep Learning Library [Internet]. arXiv; 2019 [cited 2026 May 18]. Available from: https://arxiv.org/abs/1912.01703

11. Kingma DP, Ba J. Adam: A Method for Stochastic Optimization [Internet]. arXiv; 2014 [cited 2026 May 18]. Available from: https://arxiv.org/abs/1412.6980

12. Newman AM, Liu CL, Green MR, Gentles AJ, Feng W, Xu Y, et al. Robust enumeration of cell subsets from tissue expression profiles. Nat Methods. 2015;12:453–7.

13. Benjamini Y, Hochberg Y. Controlling the False Discovery Rate: A Practical and Powerful Approach to Multiple Testing. J R Stat Soc Ser B Stat Methodol. 1995;57:289–300.

14. Ayers M, Lunceford J, Nebozhyn M, Murphy E, Loboda A, Kaufman DR, et al. IFN-γ-related mRNA profile predicts clinical response to PD-1 blockade. J Clin Invest. 2017;127:2930–40.

15. Yoshihara K, Shahmoradgoli M, Martínez E, Vegesna R, Kim H, Torres-Garcia W, et al. Inferring tumour purity and stromal and immune cell admixture from expression data. Nat Commun. 2013;4:2612.

16. Patro R, Duggal G, Love MI, Irizarry RA, Kingsford C. Salmon provides fast and bias-aware quantification of transcript expression. Nat Methods. 2017;14:417–9.

17. Durinck S, Spellman PT, Birney E, Huber W. Mapping identifiers for the integration of genomic datasets with the R/Bioconductor package biomaRt. Nat Protoc. 2009;4:1184–91.

18. Parker HS, Corrada Bravo H, Leek JT. Removing batch effects for prediction problems with frozen surrogate variable analysis. PeerJ. 2014;2:e561.

19. Sørlie T, Perou CM, Tibshirani R, Aas T, Geisler S, Johnsen H, et al. Gene expression patterns of breast carcinomas distinguish tumor subclasses with clinical implications. Proc Natl Acad Sci U S A. 2001;98:10869–74.

20. Langfelder P, Horvath S. WGCNA: an R package for weighted correlation network analysis. BMC Bioinformatics. 2008;9:559.

21. Soneson C, Love MI, Robinson MD. Differential analyses for RNA-seq: transcript-level estimates improve gene-level inferences. F1000Research. 2015;4:1521.

22. Friedman J, Hastie T, Tibshirani R. Regularization paths for generalized linear models via coordinate descent. J Stat Softw. 2010;33:1–22.

23. Feng J, Ren J, Yang Q, Liao L, Cui L, Gong Y, et al. Metabolic gene signature for predicting breast cancer recurrence using transcriptome analysis. Future Oncol. 2021;17:71–80.

24. Wang L, Zhu Y, Zhang N, Xian Y, Tang Y, Ye J, et al. The multiple roles of interferon regulatory factor family in health and disease. Signal Transduct Target Ther. 2024;9:282.

25. Purbey PK, Seo J, Paul MK, Iwamoto KS, Daly AE, Feng A-C, et al. Opposing tumor-cell-intrinsic and –extrinsic roles of the IRF1 transcription factor in antitumor immunity. Cell Rep. 2024;43:114289.

26. Smithy JW, Moore LM, Pelekanou V, Rehman J, Gaule P, Wong PF, et al. Nuclear IRF-1 expression as a mechanism to assess “capability” to express PD-L1 and response to PD-1 therapy in metastatic melanoma. J Immunother Cancer. 2017;5:25.

27. Bhat SA, Vasi Z, Adhikari R, Gudur A, Ali A, Jiang L, et al. Ubiquitin proteasome system in immune regulation and therapeutics. Curr Opin Pharmacol. 2022;67:102310.

28. Hu H, Sun S-C. Ubiquitin signaling in immune responses. Cell Res. 2016;26:457–83.

29. Hu X, Wang J, Chu M, Liu Y, Wang Z-W, Zhu X. Emerging role of ubiquitination in the regulation of PD-1/PD-L1 in cancer immunotherapy. Mol Ther J Am Soc Gene Ther. 2021;29:908–19.

30. Liu J, Cheng Y, Zheng M, Yuan B, Wang Z, Li X, et al. Targeting the ubiquitination/deubiquitination process to regulate immune checkpoint pathways. Signal Transduct Target Ther. 2021;6:28.

31. Chan XY, Singh A, Osman N, Piva TJ. Role played by signalling pathways in overcoming BRAF inhibitor resistance in melanoma. Int J Mol Sci. 2017;18:1527.

32. Grimm J, Hufnagel A, Wobser M, Borst A, Haferkamp S, Houben R, et al. BRAF inhibition causes resilience of melanoma cell lines by inducing the secretion of FGF1. Oncogenesis. 2018;7:71.

33. Konieczkowski DJ, Johannessen CM, Abudayyeh O, Kim JW, Cooper ZA, Piris A, et al. A melanoma cell state distinction influences sensitivity to MAPK pathway inhibitors. Cancer Discov. 2014;4:816–27.

34. Proietti I, Skroza N, Michelini S, Mambrin A, Balduzzi V, Bernardini N, et al. BRAF inhibitors: molecular targeting and immunomodulatory actions. Cancers. 2020;12:1823.

